# Intestinal helminth infection impairs vaccine-induced T cell responses and protection against SARS-CoV-2

**DOI:** 10.1101/2024.01.14.575588

**Authors:** Pritesh Desai, Courtney E. Karl, Baoling Ying, Chieh-Yu Liang, Tamara Garcia-Salum, Ana Carolina Santana, Felipe Ten Caten, Joseph F. Urban, Sayda M. Elbashir, Darin K. Edwards, Susan P. Ribeiro, Larissa B. Thackray, Rafick P. Sekaly, Michael S. Diamond

**Affiliations:** Department of Medicine, Washington University in St. Louis, School of Medicine, St. Louis, MO, USA; Department of Molecular Microbiology, Washington University in St. Louis, School of Medicine, St. Louis, MO, USA; Department of Pathology and Immunology, Washington University in St. Louis, School of Medicine, St. Louis, MO, USA; The Andrew M. and Jane M. Bursky Center for Human Immunology and Immunotherapy Programs, Washington University in St. Louis, School of Medicine, St. Louis, MO, USA; US Department of Agriculture, Agricultural Research Services, Beltsville Human Nutrition Research Center, Diet, Genomics, and Immunology Laboratory, and Beltsville Agricultural Research Center, Animal Parasitic Diseases Laboratory, Beltsville, MD 20705-2350, USA; Moderna, Inc., Cambridge, MA, USA; Department of Pathology and Laboratory Medicine, Emory University School of Medicine, Atlanta, GA, USA; Emory Vaccine Center, Emory University School of Medicine, Atlanta, GA, USA; Winship Cancer Institute, Emory University School of Medicine, Atlanta, GA, USA

## Abstract

Although vaccines have reduced COVID-19 disease burden, their efficacy in helminth infection endemic areas is not well characterized. We evaluated the impact of infection by *Heligmosomoides polygyrus bakeri* (Hpb), a murine intestinal hookworm, on the efficacy of an mRNA vaccine targeting the Wuhan-1 spike protein of SARS-CoV-2. Although immunization generated similar B cell responses in Hpb-infected and uninfected mice, polyfunctional CD4^+^ and CD8^+^ T cell responses were markedly reduced in Hpb-infected mice. Hpb-infected and mRNA vaccinated mice were protected against the ancestral SARS-CoV-2 strain WA1/2020, but control of lung infection was diminished against an Omicron variant compared to animals immunized without Hpb infection. Helminth mediated suppression of spike-specific CD8^+^ T cell responses occurred independently of STAT6 signaling, whereas blockade of IL-10 rescued vaccine-induced CD8^+^ T cell responses. In mice, intestinal helminth infection impairs vaccine induced T cell responses via an IL-10 pathway and compromises protection against antigenically shifted SARS-CoV-2 variants.

## INTRODUCTION

Severe acute respiratory syndrome coronavirus 2 (SARS-CoV-2), the causative agent of COVID-19 continues to threaten public health. In response to the pandemic, multiple vaccines were rapidly developed, tested, and deployed. Massive campaigns have administered more than 13.4 billion doses of vaccines corresponding to multiple platforms including mRNA, viral-vectored, protein nanoparticles, and inactivated virus vaccines (https://coronavirus.jhu.edu/map.html). Many of these vaccines had high efficacy rates early in the pandemic against symptomatic COVID-19^1^, as vaccine-induced antibody responses, particularly neutralizing antibody levels, correlated with protection^1,2^. Nonetheless, vaccine-induced T cell responses also are thought to modulate disease severity, especially in the context of SARS-CoV-2 variants that escape antibody-mediated neutralization^3–6^.

Soil-transmitted helminths that reside in the gastrointestinal (GI) tract infect more than a quarter of the global population, mostly in tropical and sub-tropical regions^7^. Most human helminth infections are caused by the hookworms *Necator americanus*, *Ancylostoma duodenale* and *Ancylostoma ceylanicum*, the roundworm *Ascaris lumbricoides,* and the whipworm *Trichuris trichiura*^8^. In healthy individuals, infection is largely asymptomatic, and adult worms can persist in the GI tract for years without causing overt pathology. However, in children and immunocompromised individuals, helminth infection in the GI tract can cause substantial morbidity. Helminth infections can skew local and systemic immune responses by eliciting a type 2 and regulatory immune response^9,10^. Although such responses can attenuate autoimmune and allergic diseases, they also can suppress clearance of pathogens that require type 1 immune response for control^11,12^.

Immune responses to vaccines vary among people living in different geographical regions^13,14^, and vaccines against tuberculosis, malaria, yellow fever, and rotavirus are less effective in Africa and Asia than in North America and Europe^15–18^. Two recent meta-analyses suggested that immune responses to vaccines against tuberculosis, measles, hepatitis B, and influenza are negatively impacted by helminth infection and that, thymus-dependent and live-virus vaccines are more likely affected^19,20^. T cell responses to Bacillus Calmette-Guerin (BCG) are reduced in people infected with helminths, and treatment with anti-helminthic drugs restores T cell proliferation and cytokine responses^21,22^. Similarly, chronic GI tract helminth infections are associated with T cell hypo-responsiveness to malaria parasites and influenza virus antigens^23^. Observations in humans are supported by animal studies, which also show detrimental effects of GI tract helminths on vaccine responses. Mice infected with schistosomes showed reduced vaccine-induced antibody responses to BCG, human papilloma virus (HPV), hepatitis B virus (HBV) and human immunodeficiency virus (HIV)^24–27^, and mice infected with the filarial helminth *Litomosoides sigmodontis* had decreased antibody responses to an influenza virus vaccine^28^. Similarly, mice infected with the intestinal hookworm *Heligmosomoides polygyrus bakeri* (Hpb) showed reduced malaria-specific antibody responses after vaccination, which correlated reduced protection against parasite challenge^29^. However, the effects of helminth infection in the GI tract on COVID-19 vaccine efficacy in humans remain unclear^30^.

Here, to begin to evaluate the impact of GI tract helminth infection on COVID-19 vaccine responses, we infected mice with GI tract-restricted Hpb and then vaccinated them with mRNA S6P, a preclinical lipid-encapsulated mRNA vaccine encoding a hexaproline stabilized Wuhan-1 spike protein of SARS-CoV-2. All Hpb-infected mice generated relatively similar anti-spike and anti-RBD specific antibody and B cell responses compared to uninfected mice. However, spike-specific CD4^+^ and CD8^+^ T cell responses were markedly decreased in Hpb-infected mice after mRNA vaccination, and similar effects were observed with an approved adenoviral-vectored (Ad26COV2.S) vaccine. Although Hpb infection during mRNA S6P priming did not impact protection against the ancestral WA1/2020 D614G virus, control of infection and disease in the lungs by the antigenically shifted BA.5.5 Omicron was diminished. The reduction of spike-specific CD8^+^ T cells after Hpb infection occurred independently of STAT6 signaling. Instead, Hpb infection induced systemic expression of IL-10, which suppressed the generation of spike-specific IFNγ^+^ CD8^+^ T cells, an effect that was reversed by blockade of IL-10 signaling. Thus, in mice, helminth infection in the GI tract inhibits systemic T cell immunity in an IL-10 dependent manner and impairs COVID-19 vaccine protection in the lung. This immune defect likely becomes important in the context of infection by SARS-CoV-2 variants that escape neutralizing antibody responses.

## RESULTS

### Intestinal helminth infection has limited effects on B cell responses to an mRNA vaccine against SARS-CoV-2

To assess whether the immunogenicity of an mRNA vaccine is affected by helminth infection, groups of 8-week-old C57BL/6J mice were primed and boosted 3 weeks later with 0.5 μg of mRNA S6P (encoding a hexaproline-stabilized Wuhan-1 (Wu-1) spike) administered by an intramuscular route. Infective Hpb (larval stage 3, L3), a GI tract-restricted helminth, was inoculated via oral gavage 12 days before prime and 12 days before booster dose (P/B) to observe the maximum potential effects on immunomodulation. Two other groups received Hpb either 12 days before the prime dose (P) or 12 days before the booster dose (B) to determine how the timing of Hpb infection impacts vaccine responses (**Fig S1A**). We confirmed Hpb infection by evaluating egg burden in feces one day before immunization with the first and/or second vaccine dose (corresponding to 11 days after Hpb inoculation) (**Fig S1B-C**). Serum samples were collected 15 days after the first or second vaccine dose, and IgG binding to Wu-1 spike protein was measured by ELISA (**Fig S1D**). Mice immunized with mRNA S6P developed spike and receptor binding domain (RBD)-specific IgG by day 15 after the first immunization, and this was enhanced at 15 days after the second vaccine dose (**Fig S1D-E**). Groups infected with Hpb (P/B, P, and B) showed similar IgG responses to spike and RBD at day 15 post-prime dose (**Fig S1D**). At day 15 post-boost, the group infected with Hpb before boost (B) showed small but significant reductions (4 to 5-fold, *P* < 0.05) in IgG responses to spike and RBD, whereas the group infected with Hpb during both prime and boost similarly reduced IgG responses (5-fold, *P* < 0.05) to RBD compared to the uninfected, mRNA S6P vaccinated group (**Fig S1E**).

We next evaluated whether Hpb infection affects vaccine-induced antigen-specific B cell responses in the spleen by interrogating germinal center (GCB; CD19^+^IgD^lo^Fas^+^GL7^+^) and memory (MBC; CD19^+^IgD^lo^Fas^-^GL7^-^TACI^-^CD138^-^) B cell subsets for binding to Wu-1 spike (**Fig S1F**). We observed no differences in the percentage or numbers of spike-specific B cell responses in Hpb-infected and uninfected mice that were vaccinated with mRNA S6P (**Fig S1G-I**). We also tested serum samples collected at day 30 post-boost to evaluate for effects of Hpb infection on neutralizing antibody responses. All vaccinated groups had similar neutralizing antibody titers against WA1/2020 D614G virus irrespective of Hpb infection (**Fig S1J**). In comparison, serum from all groups of mRNA S6P immunized mice had little to no inhibitory activity against BA.1 or BA.5 Omicron variants (**Fig S1K-L**), as reported with human serum after immunization with prototype Wu-1 spike targeted mRNA vaccines^31^. Collectively, these results demonstrate that infection with the intestinal helminth Hpb has small or no significant effects on antigen-specific B cell or neutralizing antibody responses after mRNA vaccination.

### Intestinal helminth infection impairs mRNA vaccine induced CD8^+^ T cell responses

Vaccines against SARS-CoV-2 have been shown to induce CD8^+^ T cell responses that contribute to protection^3–6^. Hence, we next investigated whether intestinal helminth infection affected mRNA vaccine induced CD8^+^ T cell responses. At day 15 post mRNA S6P boost, spike-specific IFNγ^+^TNFα^+^ CD8^+^ T cells were readily detectable in the spleen after *ex vivo* restimulation with an immunodominant spike peptide, AAAYYVGYL (spike _262-270_)^32^ (**Fig 1A-C**). However, mRNA S6P immunized mice infected with Hpb had markedly reduced IFNγ^+^TNFα^+^ CD8^+^ T cell responses. The blunted CD8^+^ T cell cytokine response in Hpb infected mice was observed regardless of whether mice were inoculated with Hpb before the first or second vaccine dose (**Fig 1A-C**).

**Figure 1.**
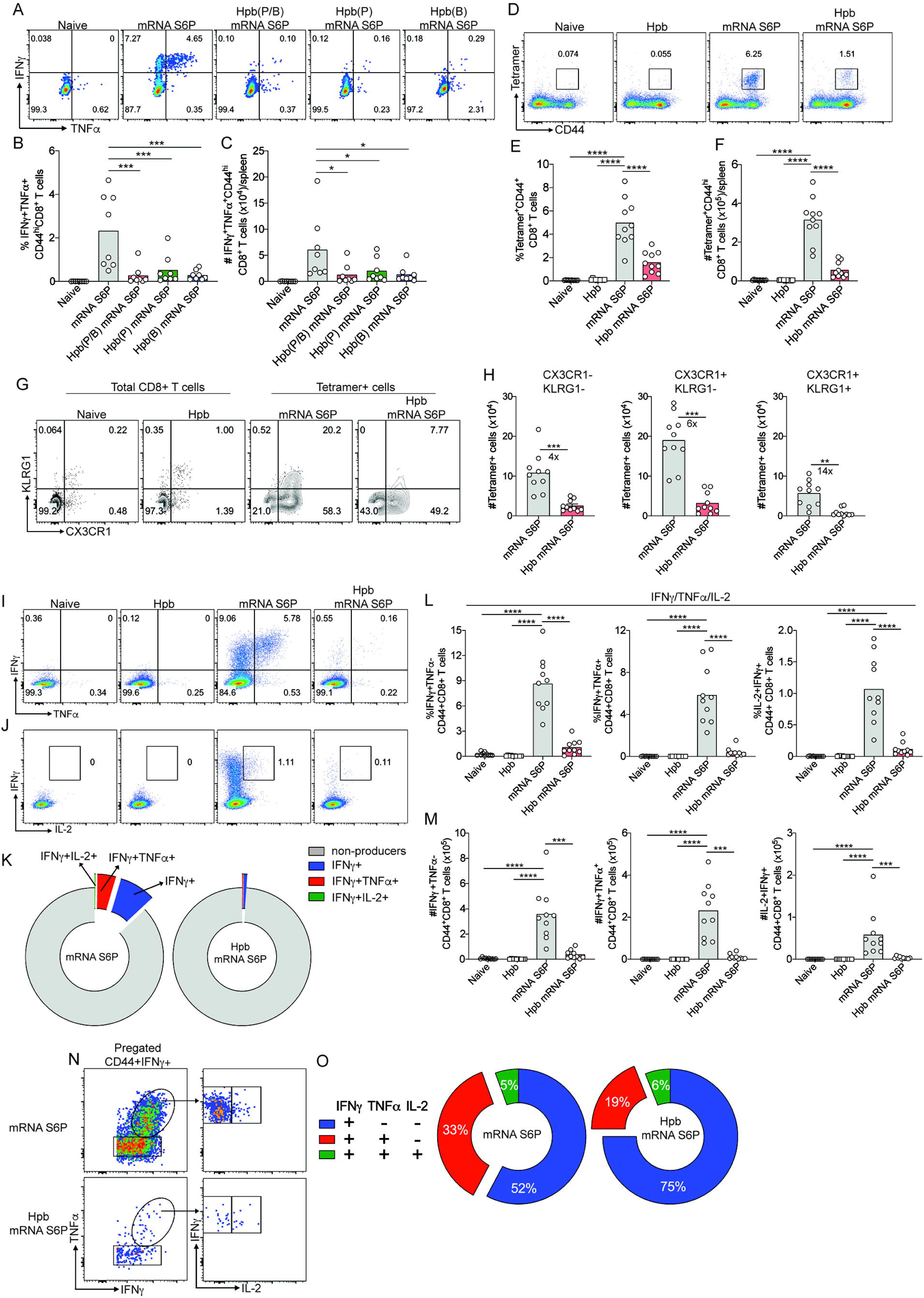
Effect of Hpb infection on CD8^+^ T cell responses in mice immunized with mRNA S6P vaccine. (**A-B**) Cohorts of C57BL/6J mice were immunized with mRNA S6P, and Hpb was gavaged either prior to prime and boost (P/B), prior to prime only (P), or prior to boost only (B). Splenocytes were harvested at day 15 post-boost, stimulated *ex vivo* with spike peptide AAAYYVGYL (S_262-270_), stained intracellularly for IFNγ and TNFα, and analyzed by flow cytometry. (**A**) Representative flow cytometry plots and the (**B**) percentages and (**C**) numbers of IFNγ and TNFα double-positive cells. (**D-O**) Cohorts of C57BL/6J mice were kept naïve, infected with Hpb alone, immunized with mRNA S6P alone, or infected with Hpb before the priming immunization with mRNA S6P. Splenocytes were isolated at day 15 post-boost, stained with antibodies and the VL8 class I MHC tetramer, and the percentage (**E**) and numbers (**F**) of tetramer-positive cells were quantified. (**G-H**) Tetramer-positive cells were analyzed for expression of CX3CR1 and KLRG1, and numbers of three subsets (CX3CR1^-^KLRG1^-^, CX3CR1^+^KLRG1^-^ and CX3CR1^+^KLRG1^+^ cells) were quantified. For naïve and Hpb-only infected mice, cells were pre-gated on total CD8^+^ T cells. (**I-O**) At day 15 post-boost, splenocytes were stimulated with spike peptide VNFNFNGL (S_539-546_), stained intracellularly for IFNγ, TNFα, and IL-2 and analyzed. Representative flow plots (**I-J**), percentages (**L**), and numbers (**M**) of IFNγ^+^TNFα^-^, IFNγ^+^TNFα^+^ and IFNγ^+^IL-2^+^ were quantified and shown as a pie charts (**K**). Non-producers are CD44^+^ cells that are negative for all three cytokines. (**N-O**) The proportionality of polyfunctional response was quantified among CD44^+^IFNγ^+^ cells and represented as pie charts. The percentages represent the mean value of the respective subset. Gating strategy is shown in **Fig S2A-B**. (**A**-**O**) Two experiments, n = 8 to 10 mice per group. Statistical analysis: (**A-F and I-M**) one-way ANOVA with Holm-Sidak’s post-test; comparisons are between uninfected and Hpb-infected mRNA S6P immunized groups or between unvaccinated and mRNA-S6P immunized groups; (**G-H**) Mann-Whitney test (\**P* < 0.05, \*\**P* < 0.01, \*\*\**P* < 0.001).

These observations prompted us to perform a more comprehensive analysis of vaccine-induced CD8^+^ T cells in mice that received two doses of Hpb (P/B). Here, we identified spike-specific CD8^+^ T cells by staining with H-2K^b^ restricted immunodominant class I MHC tetramers (VL8, VNFNFNGL: spike _539-546_)^32^. Uninfected, vaccinated mice had approximately 5% tetramer-positive CD8^+^ T cells on day 15 post-boost in the spleen. In comparison, we observed reduced frequency (3-fold lower, *P* < 0.0001) and number (6-fold lower, *P* < 0.0001) of tetramer-positive CD8^+^ T cells in Hpb-infected mice (**Fig 1D-F**). As expected, naïve mice or mice infected with Hpb alone did not generate spike-specific CD8^+^ T cells. We assessed whether Hpb infection preferentially impacted a subset of effector and memory CD8^+^ T cells using the phenotypic markers CX3CR1 and KLRG1^33^. CD8^+^ T cells from naïve and Hpb-only infected mice expressed little or no CX3CR1 or KLRG1 (**Fig 1G**). However, in mRNA S6P vaccinated mice, most tetramer-positive cells expressed CX3CR1, 20% of which also expressed KLRG1, representing terminally differentiated effector cells (**Fig 1G-H**). In contrast, only 8% of tetramer-positive CD8^+^ T cells in Hpb-infected mice immunized with mRNA S6P were CX3CR1^+^KLRG1^+^, and this corresponded to 14-fold (*P* < 0.01) lower numbers compared to uninfected, vaccinated mice (**Fig 1G-H**). The other two subsets CX3CR1^-^KLRG1^-^ and CX3CR1^+^KLRG1^-^ that represent memory precursors and less differentiated effector cells respectively, were reduced by approximately 4 to 6-fold (*P* < 0.001) in vaccinated Hpb-infected mice compared to vaccinated uninfected mice.

We performed additional functional analysis by restimulating CD8^+^ T cells *ex vivo* with the VL8 spike peptide and measuring intracellular cytokine levels. Vaccinated mice infected with Hpb showed decreases in both the percentages and total numbers of IFNγ^+^TNFα^-^, IFNγ^+^TNFα^+^ and IL-2^+^IFNγ^+^ CD8^+^ T cells (**Fig 1I-M, S2A**), suggesting that CD8^+^ T cell effector responses are suppressed due to Hpb infection. To assess effects on the polyfunctional response, we evaluated the percentage of cells among CD44^+^IFNγ^+^ cells that co-express TNFα and/or IL-2 (**Fig S2B**). Among CD44^+^ IFNγ^+^ CD8^+^ T cells, Hpb-infected mice immunized with mRNA S6P had reduced proportions of TNFα^+^ but not IL-2^+^ cells (**Fig 1N-O**) compared to uninfected, vaccinated mice. We also assessed whether Hpb infection polarized vaccine-induced CD8^+^ T cells towards type 2 and regulatory phenotypes by measuring IL-4 and IL-10 responses after *ex vivo* VL8 spike peptide restimulation. Vaccinated mice infected with Hpb had similar percentages and numbers of IL-4^+^ and IL-10^+^ cells after spike peptide exposure compared to uninfected vaccinated mice, and these levels were not greater than the background response observed in naïve or Hpb-infected mice (**Fig S2C-F**). Together, these results show that Hpb infection suppresses priming, differentiation, and polyfunctionality of mRNA S6P vaccine-induced CD8^+^ T cells but does not induce production of type 2 or regulatory cytokines.

### Intestinal helminth infection impairs vaccine-induced CD4^+^ T_H_1 T cell responses

We next examined whether Hpb infection also causes defects in CD4^+^ T cell responses after immunization with the mRNA S6P vaccine. At day 15 post-boost, uninfected mice immunized with mRNA S6P had detectable IFNγ^+^TNFα^+^ CD4^+^ T cells after *ex vivo* stimulation of splenocytes with an immunodominant class II MHC restricted peptide VTWFHAIHVSGTNGT (spike _62-76_)^32^ (**Fig 2A**). In contrast, Hpb infection resulted in impaired IFNγ^+^TNFα^+^ CD4^+^ T responses (**Fig 2A-C**). Although all three Hpb-infected groups (P, B, and P/B) showed reduced fractions of CD4^+^ T cells expressing IFNγ and TNFα, only those receiving Hpb prior to both immunizations (P/B) showed reduced numbers (**Fig 2A-C**). Given this result, in subsequent analysis of CD4^+^ T cell responses, we focused on the P/B group. The overall percentages and numbers of IFNγ^+^ TNFα^-^, IFNγ^+^TNFα^+^ and IL-2^+^IFNγ^+^ CD4^+^ T cells were reduced in Hpb-infected compared to uninfected vaccinated mice (**Fig 2F-G and S3A**). In terms of polyfunctionality of CD44^+^IFNγ^+^ CD4^+^ T cells, Hpb-infected mice immunized with mRNA S6P showed reduced proportions of TNFα^+^ cells (**Fig 2D-E**). However, intracellular levels of IL-4 or IL-10 in spike peptide restimulated CD4^+^ T cells from vaccinated mice infected with Hpb were equivalent to those seen in naïve or uninfected, vaccinated mice (**Fig 2H-I and S3B**). We also assessed T_H_2 responses by staining for the canonical transcription factor GATA-3 to determine if helminth infection skews the differentiation profile of CD4^+^ T cells induced by mRNA S6P vaccination. Mice infected with Hpb with or without mRNA S6P vaccination had higher levels of T_H_2 (Foxp3^-^ GATA-3^+^CD44^+^) cells in the spleen than uninfected mice. Thus, helminth infection skews towards T_H_2 differentiation, and vaccination does not affect this response (**Fig S3C-D**).

**Figure 2.**
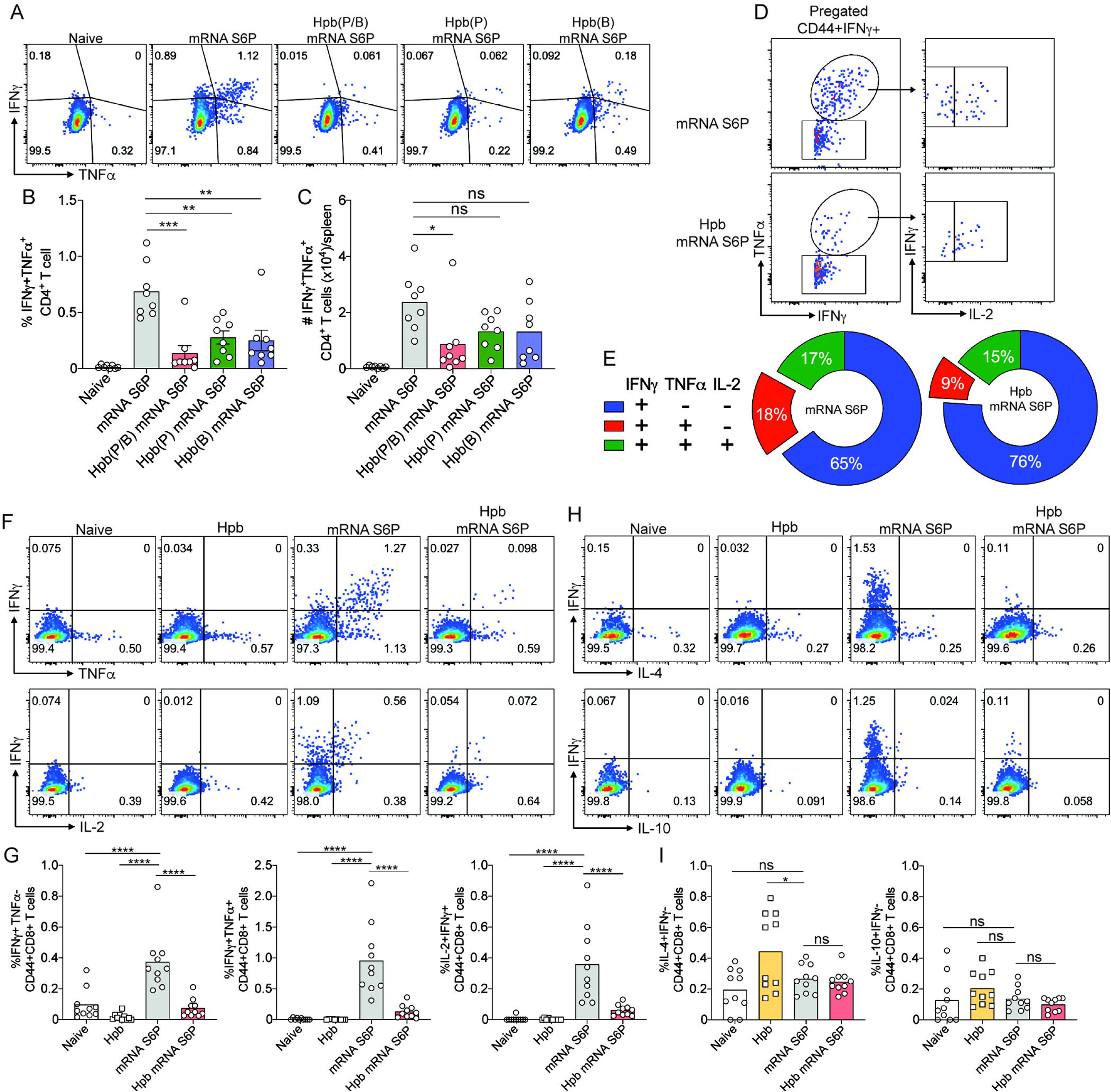
Effect of Hpb infection on CD4^+^ T cell responses in mice immunized with mRNA S6P vaccine. (**A-B**) Cohorts of C57BL/6J mice were immunized with mRNA S6P, and Hpb was given either prior to prime and boost (P/B), prior to prime (P), or prior to boost (B). Splenocytes were stimulated *ex vivo* with spike peptide VTWFHAIHVSGTNGT (S_62-76_), stained intracellularly for IFNγ and TNFα, and analyzed by flow cytometry. (**A**) Representative flow cytometry plots, (**B**) percentages, and (**C**) numbers of IFNγ^+^TNFα^+^ cells. (**D-I**) Cohorts of C57BL/6J mice were immunized and treated with Hpb only before vaccine priming. (**D-G**) At day 15 post-boost, splenocytes were stimulated with spike peptide, stained intracellularly for IFNγ, TNFα and IL-2 and analyzed. (**D**-**E**) Representative flow cytometry plots of IFNγ^+^TNFα^+^ and IFNγ^+^TNFα^+^IL-2^+^ cells are shown. Polyfunctional responses were quantified among CD44^+^IFNγ^+^ CD4^+^ T cells and are represented as pie charts. (**F-G**) Representative flow cytometry plots and percentages of IFNγ^+^ single-positive and IFNγ^+^TNFα^+^ and IL-2^+^IFNγ^+^ double-positive cells are shown. (**H-I**) After peptide stimulation, splenocytes also were stained intracellularly for IFNγ, IL-4 and IL-10. Representative flow cytometry plots and percentages of IL-4 positive and IL-10 positive cells are shown. Two experiments, n = 8 to 10 mice per group. Statistical analysis: one-way ANOVA with Holm-Sidak’s post-test; comparisons are between uninfected and Hpb-infected mRNA S6P immunized groups or between unvaccinated and S6P-mRNA immunized groups (ns, not significant, \**P* < 0.05, \*\**P* < 0.01, \*\*\**P* < 0.001, \*\*\*\**P* < 0.0001).

### Intestinal helminth infection impairs adenoviral vectored vaccine-induced T cell response

We next assessed the generalizability of these findings by evaluating responses to an adenoviral-vectored vaccine (Ad26.COV2.S; Janssen) that was deployed in humans and induced robust T cell responses^34,35^. Twelve days after gavaging C57BL/6 mice with Hpb, we immunized animals by intramuscular injection with 5 x 10^9^ viral particles of Ad26.COV2.S and then boosted them 30 days later (**Fig S4A**). Ten days after boosting, we observed that while the percentages of tetramer-positive CD8^+^ T cells were unaffected by infection with Hpb, there was an approximately 2-fold decrease (*P* < 0.01) in the number of spike-specific CD8^+^ T cells (**Fig S4B-C**). Moreover, vaccinated mice infected with Hpb had reduced proportions and numbers of CX3CR1^+^KLRG1^+^ terminally differentiated effector CD8^+^ T cells compared to uninfected mice immunized with Ad26.COV2.S (**Fig S4D-E**). In functional studies after restimulation with the class I MHC-restricted immunodominant peptide VL8, Hpb infected mice vaccinated with Ad26.COV2.S had reduced percentages and numbers of IFNγ^+^ TNFα^−^, IFNγ^+^ TNFα^+^, and IFNγ^+^ IL-2^+^ CD8^+^ T cells than uninfected and immunized mice (**Fig S4F-G**). Similar to the results with the mRNA S6P vaccine, skewing of the CD8^+^ T cell response to IL-4 or IL-10 after peptide restimulation was not observed in Hpb-infected, Ad26.COV2.S-vaccinated mice (**Fig S4H-I**).

We also assessed if Hpb infection induced defects in the spike-specific CD4^+^ T cell response after immunization with Ad26.COV2.S. Indeed, infection with Hpb resulted in decreased percentages and numbers of IFNγ^+^ CD4^+^ T cells after peptide restimulation (**Fig S4J-K**). While the percentage of IFNγ^+^TNFα^+^ CD4^+^ T cells also was reduced in Hpb-infected Ad26.COV2.S-vaccinated mice, the numbers were not. Hpb infection also did not skew the Ad26.COV2.S vaccine-induced CD4^+^ T cell response to producing IL-4 or IL-10 **(Fig S4L and M**). Collectively, these results show that Hpb infection also impairs Ad26.COV2.S vaccine induced CD8^+^ and CD4^+^ T cell responses, although to a lesser degree than that observed with the mRNA S6P vaccine.

### Intestinal helminth infection impairs vaccine protection against a SARS-CoV-2 Omicron variant

We investigated whether the defects in immune responses associated with Hpb infection during mRNA vaccination would compromise protection against SARS-CoV-2. Groups of 6-week-old K18-hACE2 mice were gavaged with Hpb, and 12 days later were immunized and subsequently boosted with mRNA S6P vaccine (**Fig S5A**). Naïve K18-hACE2 mice and Hpb-infected unvaccinated mice served as controls. We first assessed helminth infection in K18-hACE2 mice and observed comparable egg burdens as seen in C57BL6/J mice (**Fig S5B**). We also confirmed that Hpb infection before the prime dose negatively affects antigen-specific CD8^+^ T cell responses in K18-hACE2 mice (**Fig S5C**).

Four to 5 weeks after boosting, mice were challenged by intranasal route with 10^4^ focus-forming units (FFU) of WA1/2020 D614G or an Omicron variant, BA.5.5. After WA1/2020 D614G infection, all unvaccinated mice (naïve or Hpb-infected) showed weight loss on days 4-and 5 post-virus infection (dpi) (**Fig 3A**). In contrast, all vaccinated mice irrespective of Hpb infection were protected from weight loss. Compared to unvaccinated (naïve or Hpb-infected only) mice, animals immunized with mRNA S6P had lower viral burden at 5 dpi in the nasal wash, nasal turbinate, and lung (**Fig 3B-E**). Vaccinated mice that had been infected with Hpb also had reduced viral burdens, suggesting that helminth infection did not impact protection against WA1/2020 D614G (**Fig 3B-E**), possibly due to the equivalent serum neutralizing activity against WA1/2020 D614G in the presence or absence of Hpb (**Fig S1J**). In contrast, all mRNA S6P vaccinated groups showed poor serum neutralizing antibody responses against BA.5 (**Fig S1L**). Following challenge with BA.5.5, unvaccinated and Hpb-infected vaccinated mice lost 15% of body weight over a 5-day period, whereas vaccinated-only mice maintained their body weight (**Fig 3F**). At 5 dpi, unvaccinated mice had higher levels of BA.5.5 RNA in nasal washes and nasal turbinates than mRNA S6P vaccinated mice, but Hpb infection did not affect this protection (**Fig 3G-H**). In the lung, unvaccinated mice also had higher viral burden than mice immunized with mRNA S6P. Moreover, Hpb-infected vaccinated mice had higher levels of viral RNA or infectious virus in the lung than animals immunized with mRNA S6P and not infected with Hpb (**Fig 3I-J**). Overall, these data suggest that Hpb infection compromises vaccine-mediated mediated protection in the lung against the antigenically shifted BA.5.5 Omicron variant, possibly due to impaired T cell responses.

**Figure 3.**
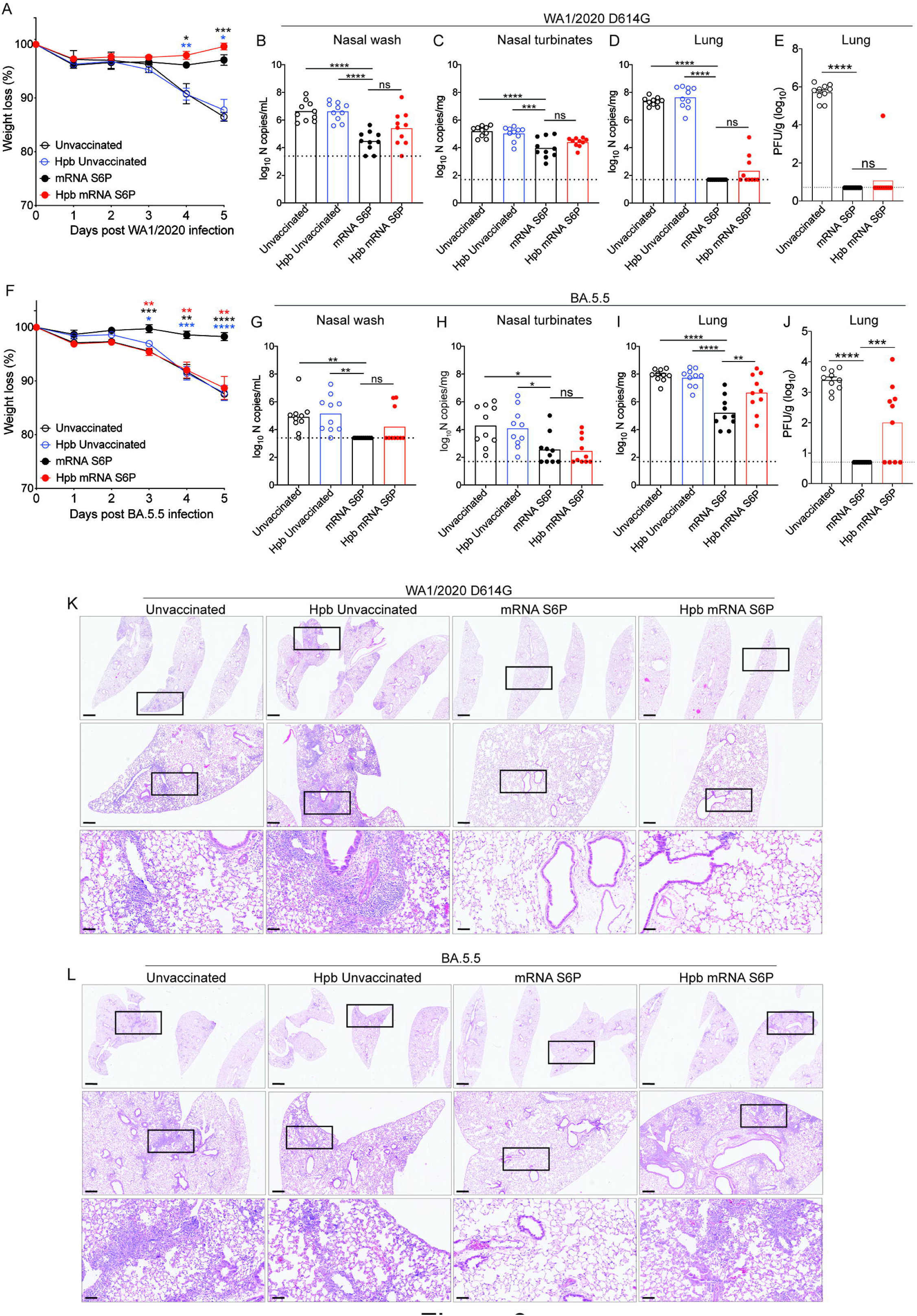
Effect of Hpb infection on mRNA S6P vaccine-mediated protection. (**A-L**) Six-week-old K18-hACE2 mice were immunized with 0.5 μg of mRNA S6P, and infective Hpb L3 was given prior to the priming dose. As controls, some mice were kept unvaccinated (naïve) or given Hpb without vaccination. All mice were challenged intranasally at day 40 post-boost (or similarly aged controls) with 10^4^ PFU of WA1/2020 D614G (**A-E**) or BA.5.5 (**F-J**). Weight was monitored through 5 dpi (**A, F**). Viral burden at 5 dpi was assessed by qRT-PCR (**B-D, G-I**) or plaque assay (**E, J**) in nasal washes (**B, G**), nasal turbinates (**C, H**), and lungs (**D, E, I, J**) after inoculation with WA1/2020 D614G (**B-E**) or BA.5.5 (**G-J**). Two experiments, n = 10 mice per group. (**K-L**) Hematoxylin and eosin staining of lung sections harvested at 5 dpi after WA1/2020/D614G (**K**) or BA.5.5 inoculation (**L**). Low (top: scale bars, 1 mm), moderate (middle: scale bars, 200 μm), and high (bottom: scale bars, 50 μm) power images are shown. Representative images of multiple lung sections from *n* = 6 from each group, two experiments. Statistical analysis: (**A, F**) two-way ANOVA with Dunnett’s post-test; comparisons are between mRNA S6P vaccinated mice and all other groups; (**B, C, D, E, G, H, I, J**) one-way Kruskal– Wallis ANOVA with Dunn’s post-test; comparisons are between mRNA S6P vaccinated mice and all other groups; dotted lines show LOD (ns, not significant, \**P* < 0.05, \*\**P* < 0.01, \*\*\**P* < 0.001, \*\*\*\**P* < 0.0001).

As an independent metric of protection, we performed histological analysis of lungs at 5 dpi after infection with WA1/2020 D614G or BA.5.5. Lungs of unvaccinated mice infected with WA1/2020 D614G showed evidence of severe pneumonia characterized by immune cell infiltration, alveolar space consolidation, and vascular congestion (**Fig 3K**). mRNA S6P vaccinated mice infected with WA1/2020 D614G were protected from lung pathology irrespective of Hpb infection (**Fig 3K**). BA.5.5 infection of unvaccinated mice caused infiltration of immune cells in the lungs with patchy alveolar space consolidation (**Fig 3L**), and this was prevented by mRNA S6P vaccination. However, Hpb-infected vaccinated mice had residual lung inflammation after BA.5.5 challenge (**Fig 3L**), consistent with the higher lung viral burden (**Fig 3I-J**). Together, these results suggest that Hpb infection impairs mRNA S6P vaccine-mediated protection against lung infection and inflammation caused by antigenically shifted BA.5.5 Omicron variants.

### Intestinal helminth infection durably affects vaccine induced CD8^+^ T cell response prior to boosting

We next evaluated if Hpb impacted vaccine induced CD8^+^ T cell responses during the priming phase, prior to boosting. We harvested spleens from naïve mice, Hpb-infected mRNA S6P vaccinated mice, and uninfected mRNA vaccinated mice at day 21 post-prime and analyzed CD8^+^ T cell responses. The percentage and number of spike-specific CD8^+^ T cells were reduced in Hpb infected vaccinated mice compared to uninfected mice receiving vaccine only (**Fig 4A and C**). Splenic CD8^+^ T cells stimulated *ex vivo* with VL8 spike peptide also had significant decreases in percentages and numbers of IFNγ^+^TNFα^−^ and IFNγ^+^TNFα^+^ cells in helminth infected mice vaccinated with mRNA S6P (**Fig 4B, D, E**).

**Figure 4.**
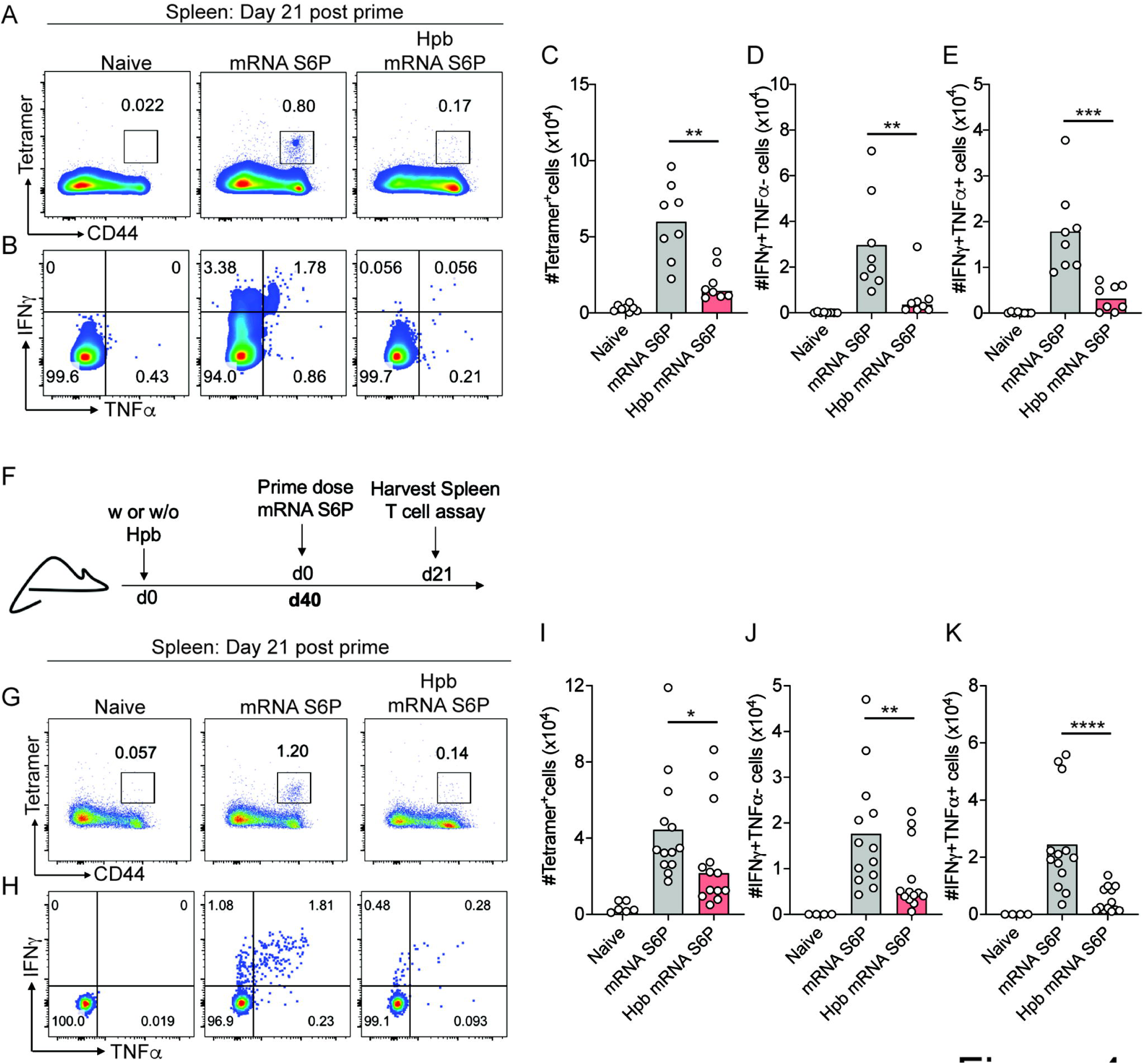
Helminth infection affects vaccine induced CD8^+^ T cell responses prior to boosting. (**A-E**) Mice were immunized once with mRNA S6P, and Hpb was administered prior to immunization. At day 21 post-prime, splenocytes were stained with VL8 tetramer, representative flow plots (**A**) and numbers (**C**) of tetramer^+^ cells are shown. (**B, D-E**) Splenocytes were stimulated with VL8 peptide and numbers of IFNγ^+^TNFα^-^ (**D**) and IFNγ^+^TNFα^+^ cells (**E**) were quantified. (**F**) Schematic representation of experimental design showing mice infected with Hpb and 40 days later primed with one dose of mRNA S6P. At 21 days post-immunization, splenocytes were stained with CD3, CD8, CD44 and VL8 tetramer. Representative flow plots (**G**) and numbers (**I**) of tetramer positive CD8^+^ T cells were quantified. (**H, J-K**) Splenocytes were stimulated with VL8 peptide and stained intracellularly with IFNγ and TNFα. Representative flow plots (**H**) and numbers of IFNγ^+^TNFα^-^ (**J**) and IFNγ^+^TNFα^+^ (**K**) were quantified. (**A-E**) Two experiments, n = 8 mice per group. (**G-K**) Three experiments, *n* = 13 mice per group. Statistical analysis: one-way ANOVA with Holm-Sidak’s post-test; comparisons are between uninfected and Hpb-infected mRNA S6P immunized groups (\**P* < 0.05, \*\**P* < 0.01, \*\*\**P* < 0.001, \*\*\*\**P* < 0.0001).

To examine whether Hpb infection affects T cell cytokine production in a spike antigen-independent manner, we stimulated splenocytes with anti-CD3 and anti-CD28 antibodies. Compared to mice vaccinated with mRNA S6P alone, helminth infected mice vaccinated with mRNA S6P had significant defects in both percentages and numbers of IFNγ^+^TNFα^-^ and IFNγ^+^TNFα^+^ CD8^+^ T cells after anti-CD3/C28 stimulation, like that seen after incubation with VL8 peptide (**Fig S6C-D**). To test whether Hpb mediated defects in T cell activity requires T cell receptor (TCR) crosslinking, we stimulated splenocytes with PMA (phorbol-12-myristate-13-acetate) and ionomycin, which bypasses the membrane-proximal steps of TCR signaling. While PMA/ionomycin robustly induced cytokine production in CD8^+^ T cells of mRNA S6P vaccinated mice, this effect was diminished in animals infected with Hpb (**Fig S6C-D**). Collectively, these studies suggest that Hpb infection renders CD8^+^ T cells less responsive to antigen-dependent and independent stimulation.

Because defects in vaccine-induced CD8^+^ T cell responses were observed at day 21 post-prime in helminth infected mice (**Fig 4A-E**), we focused on this time point in further experiments. While Hpb causes infection in C57BL/6 mice, the immunomodulatory effects and egg burden decrease after 3 weeks resulting in a lower level of chronic infection^36^. To assess the durability of the effect of Hpb infection on CD8^+^ T cell responses, we immunized mice with mRNA S6P vaccine at day 40 rather than day 12 post Hpb infection (**Fig 4F**). Even at this later time point, tetramer^+^ and IFNγ^+^TNFα^−^ and IFNγ^+^TNFα^+^ CD8^+^ T cells in the spleen were lower after peptide restimulation in Hpb infected mice than animals receiving vaccine alone (**Fig 4G-K**). These data suggest that Hpb infection causes defects in vaccine induced CD8^+^ T cell response prior to boosting that last for at least 40 days.

### STAT6 deficiency does not rescue helminth induced defects in vaccine CD8^+^ T cell responses

Helminth infection elicits type 2 immune responses, which are largely dependent on the transcription factor STAT6^37^. STAT6 signaling is a primary pathway by which enteric helminth infection hinders antiviral T cell immunity in coinfection studies^38,39^. To investigate whether STAT6 signaling mediates Hpb triggered defects in vaccine generated CD8^+^ T cell responses, we repeated Hpb infection and mRNA vaccination in *Stat6*^-/-^ and congenic WT mice. In the absence of Hpb infection, mRNA vaccination generated equivalent CD8^+^ T cell responses in WT and *Stat6*^-/-^ mice (**Fig 5A-E**). Furthermore, tetramer^+^ CD8^+^ T cell response and effector cytokine responses (IFNγ^+^TNFα^−^ and IFNγ^+^TNFα^+^) were similarly diminished in *Stat6*^-/-^ and WT mice infected with Hpb (**Fig 5A-E**). Thus, Hpb infection dampens CD8^+^ T cell responses to mRNA S6P vaccine through a STAT6-independent signaling pathway.

**Figure 5.**
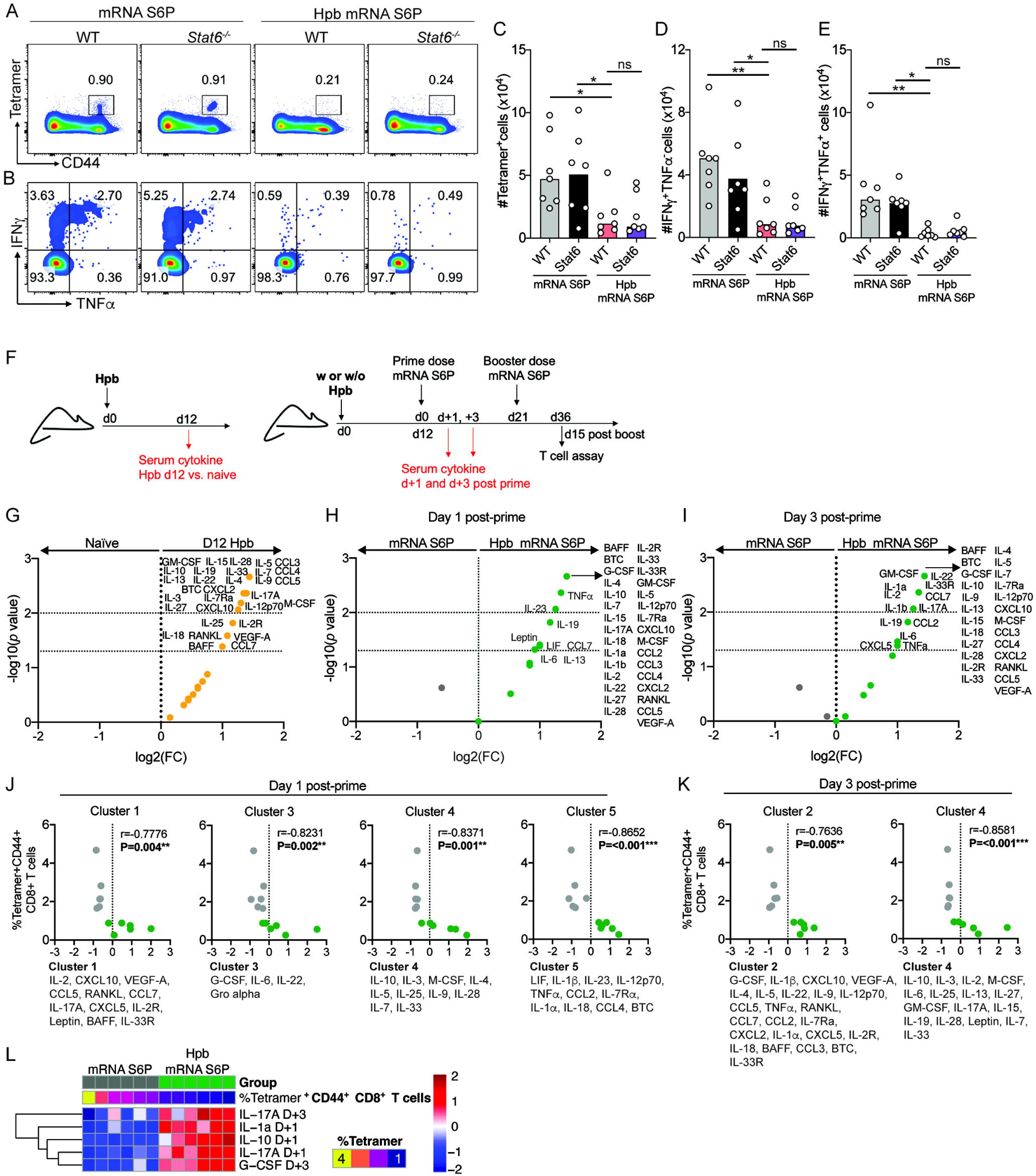
STA6-independent suppression of vaccine induced CD8^+^ T cell response by intestinal helminths. (**A-E**) WT and *Stat6*^-/-^ mice were immunized with one dose of mRNA S6P, and Hpb was administered prior to immunization. At day 21 post-prime, splenocytes were stained with VL8 tetramer, representative flow plots (**A**) and numbers (**C**) of tetramer^+^ cells are shown. (**B, D-E**) Splenocytes were stimulated with VL8 peptide and numbers of IFNγ^+^TNFα^-^ (**D**) and IFNγ^+^TNFα^+^ cells (**E**) were quantified. (**F**) Schematic representation of experimental design. Mice were immunized with mRNA S6P, and Hpb was given 12 days prior to the priming dose. (**G**) Serum samples were collected at day 12 post Hpb infection (yellow dots) or from uninfected mice, and cytokine responses were measured by Luminex assay and represented using volcano plots. (**H-L**) Serum samples were collected after day 1 (**H**) and 3 (**I**) post-prime vaccination, and cytokine response was compared between Hpb infected mRNA S6P immunized mice (green dots) and uninfected mice immunized with mRNA S6P (grey dots). (**J-K**) Spearman correlation analysis of cytokine clusters between uninfected (grey dots) and Hpb-infected (green dots) mice vaccinated with mRNA S6P at day 1 (**J**) and 3 post-prime (**K**) with the percentage of VL8 tetramer-positive CD8^+^ T cells in the spleen at day 15 post-boost. (**L**) Feature selection of serum cytokines at days 1 and 3 post-prime shown as a heatmap that relates to the percentage of VL8 tetramer-positive CD8^+^ T cells at day 15 post-boost between uninfected and Hpb-infected mRNA S6P vaccinated mice. (**A-E**) Two experiments, n = 8 mice per group. Statistical analysis: one-way ANOVA with Holm-Sidak’s post-test; comparisons are Hpb infected mRNA S6P immunized WT mice (red) to other groups (ns, not significant, \**P* < 0.05, \*\**P* < 0.01). (**G-I**) Cytokine differential expression volcano plots were generated by plotting log_2_ (fold-change) vs -log_10_ (*P* value), and the *P* values were obtained by Mann-Whitney tests with repeated measure corrections, \*\**P* < 0.01; FC, fold-change (n = 6 per group). (**J-L**) Clustering analysis of cytokines levels was performed with K-means clustering method (see Methods). For feature selection, scaled values of significantly different cytokines (*P* < 0.05) were used as input for a Lasso regression (glmnet R package -v 4.1-7), and cytokines with non-zero coefficients were selected. Nonparametric Spearman correlation analysis was performed using a 95% confidence interval (** *P* < 0.01, *** *P* < 0.001).

To explore alternate mechanisms associated with suppression of vaccine-induced T cell immune responses in Hpb-infected mice, we collected serum at early timepoints after prime immunization and profiled cytokines (**Fig 5F**). We first collected serum from mice infected with Hpb and assessed cytokine levels. As expected, several type 2 and regulatory cytokines (*e.g*., IL-4, IL-5, IL-13, IL-25, IL-33, and IL-10) were upregulated in Hpb-infected serum samples at 12 dpi compared to naïve mice (**Fig 5G**). Unexpectedly, Hpb infection also resulted in an increase of type 1 and 17 signature cytokines including IL-12p70, IL-27, IL-17A, and IL-22, and inflammatory mediators including CCL3, CCL4, CCL5, CCL7, CXCL2, and CXCL10 compared to naïve serum (**Fig 5G**). We then immunized naïve and Hpb-infected mice with mRNA S6P and evaluated serum cytokine levels at day 1 and 3 post-prime. Hpb-infected and vaccinated mice had higher serum cytokine responses than mice immunized with mRNA S6P alone (**Fig 5H-I**). Several type 1, type 2, and type 17 signature cytokines as well inflammatory mediators were higher in Hpb-infected vaccinated mice than in uninfected vaccinated mice at days 1 and 3 post-prime (**Fig 5H-I**). To further evaluate the distribution of cytokines expression in the groups, we performed unsupervised analysis using a clustering method. This analysis identified 5 optimal cytokine clusters at day 1 post-prime and 4 clusters at day 3 post-prime. At day 1 post-prime, clusters 1, 3, 4 and 5, and at day 3 post-prime, clusters 4 and 5 inversely correlated with tetramer-positive CD8^+^ T cell response in these mice (**Fig 5J-K and S7A-B**). However, these clusters did not reveal a distinct pattern for the segregation of cytokines and inflammatory mediators when comparing different conditions. Therefore, we performed feature analysis and identified IL-1α, IL-10, IL-17A, and G-CSF as having a determinant segregating role in the reduced T cell response observed under helminth pre-infection conditions (**Fig 5L**). Moreover, in Hpb-infected *Stat6*^-/-^ mice, although type 2 cytokines IL-4, IL-5, IL-13, IL-33R were downregulated, elevated levels of G-CSF, IL-17A, IL-10 and IL-1α were still present (**Fig S7C**). Thus, Hpb infection induces a mixed type 1, type 2, and type 17 cytokine response signature. Four of these cytokines negatively correlated with vaccine-induced CD8^+^ T cells, and these were not affected in Hpb-infected *Stat6*^-/-^ mice, which also had blunted CD8^+^ T cell responses after mRNA vaccination.

### Hpb causes defects in vaccine induced CD8^+^ T cell responses in an IL-10 dependent manner

We focused on IL-10 because of its role as an immunosuppressive cytokine^40^. Although helminth infection can stimulate expression of IL-10 in the GI tract, its cellular source of IL-10 in secondary lymphoid tissues is less clear^41^. To identify the cell types that produce IL-10, we infected 10BiT (IL-10) reporter mice^42^ with Hpb and then immunized them with mRNA S6P. Other cohorts included 10BiT reporter mice that received mRNA S6P alone and naïve animals. 10BiT reporter mice possess a Thy1.1 cDNA inserted upstream of the *IL10* locus and as a result, cells transcribing IL-10 express Thy1.1 on their plasma mambrane^42^. These mice also were crossed to reporter mice with sequences encoding green fluorescence protein (GFP) knocked into the *Foxp3* gene (Foxp3^gfp^ mice) to generate dual reporter mice^42^. Because we speculated that IL-10 might have an early effect during T cell priming, we analyzed its expression at 3 days after priming. In splenocytes, we observed an increased (albeit still low) frequency of IL-10 expressing CD4^+^ T cells (Foxp3^+^ and Foxp3^-^), CD8^+^ T cells, NK T cells, and NK cells in helminth infected mRNA S6P vaccinated mice than naïve or mRNA S6P immunized mice alone (**Fig 6A-B and S8**). A subset of CD8^+^ T cells, NK T cells, and NK cells from Hpb-infected animals produced more IL-10 than their uninfected control counterparts. Although there was a trend towards higher IL-10 expressing cells among B cells and DC subsets in helminth infected mice, this did not attain statistical significance. The pattern of IL-10 expression in cytolytic lymphocytes is consistent with the elevated levels of IL-10 in serum in Hpb-infected mice.

**Figure 6.**
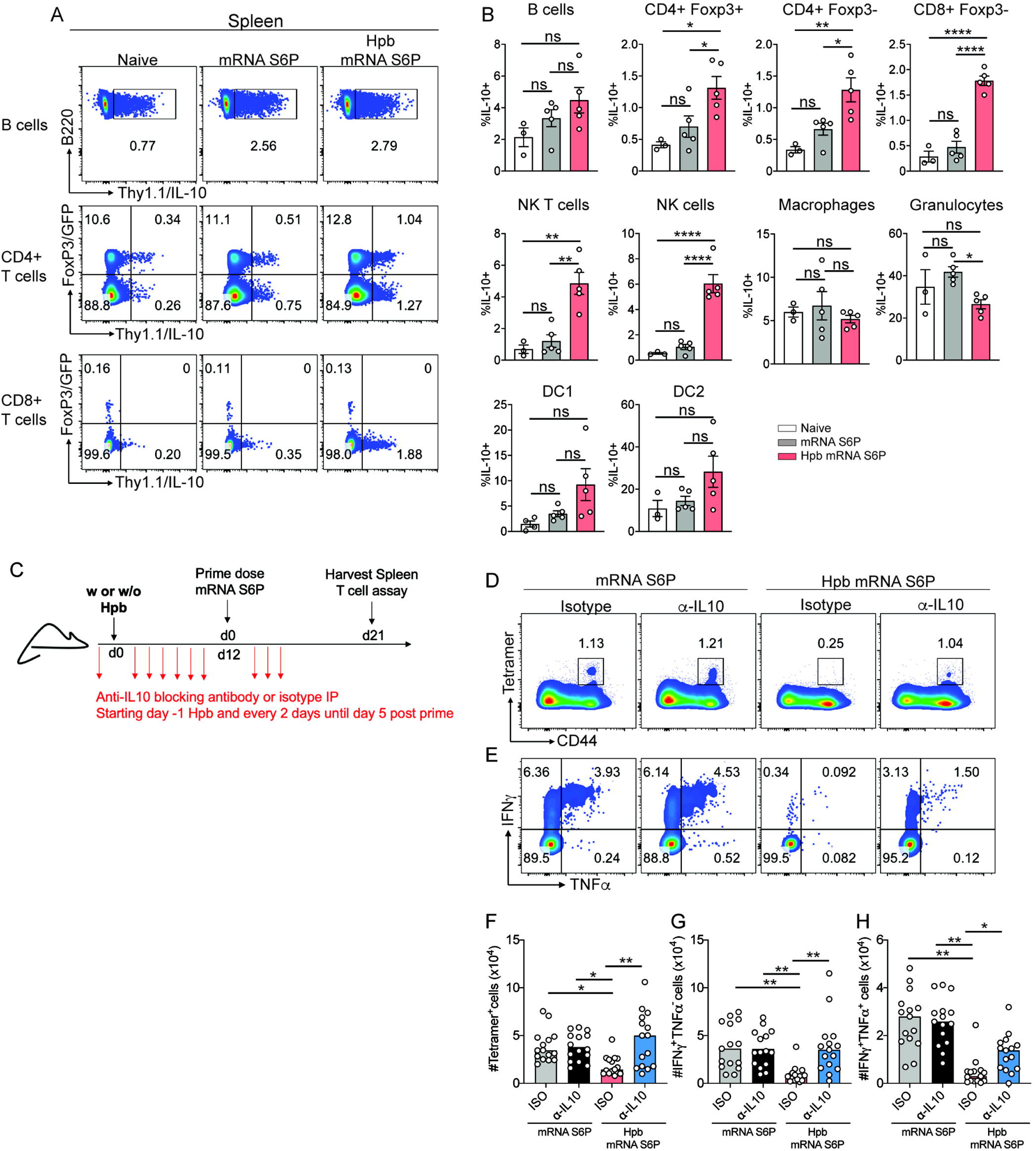
Blockade of IL-10 in helminth infected mice restores CD8^+^ T cell responses to mRNA S6P. (**A-B**) IL-10-Thy1.1 10BiT IL-10 reporter/Foxp3-GFP reporter mice were infected with Hpb and immunized with one dose of mRNA S6P. Three days post prime, splenocytes were stained with various cell surface immune cell markers and anti-Thy1.1 for IL-10 expression. (**A**) Representative flow plots of B cells, CD4^+^ T cells (Foxp3^+^ and Foxp3^-^) and CD8^+^ T cells are shown. (**B**) Percentages of IL-10 expressing cells among B cells, CD4^+^Foxp3^+^ T cells, CD4^+^Foxp3^-^ T cells, CD8^+^ T cells, NK T cells, NK cells, macrophages, granulocytes, type 1 and type 2 dendritic cells (DC1 and DC2) were quantified. (**C-H**) Hpb infected mice immunized with mRNA S6P received anti-IL10 blocking antibody or isotype every two days, starting one day prior to Hpb infection, and continued until day 5 post-prime vaccination. (**C**) Schematic representation of experimental design. At day 21 post-prime, splenocytes were stained with VL8 tetramer, representative flow plots (**D**) and numbers (**F**) of tetramer^+^ cells are shown. (**E, G-H**) Splenocytes were stimulated with VL8 peptide and numbers of IFNγ^+^TNFα^-^ (**G**) and IFNγ^+^TNFα^+^ cells (**H**) were quantified. (**A-B**) Two experiments, n = 3 to 5 mice per group. Statistical analysis: one-way ANOVA with Holm-Sidak’s post-test; all three groups are compared with each other (ns = not significant, \**P* < 0.05, \*\**P* < 0.01, \*\*\*\**P* < 0.0001). (**F-H**) Three experiments, n = 15 mice per group. Statistical analysis: one-way ANOVA with Holm-Sidak’s post-test; comparisons are between Hpb infected mRNA S6P immunized mice that received isotype antibody (red) to all other groups (ns, not significant, \**P* < 0.05, \*\**P* < 0.01, \*\*\**P* < 0.001).

To test whether the suppression of vaccine-induced T cell responses in helminth infected mice is mediated by IL-10, we blocked its function by injecting anti-IL-10 antibody starting one day prior to Hpb infection and administering it every 2 days until 5 days after the priming dose of mRNA S6P vaccine (**Fig 6C**). Blockade of IL-10 did not enhance CD8^+^ T cell responses in mice receiving mRNA vaccine alone (**Fig 6D-H**). However, anti-IL-10 treatment of helminth infected mice restored tetramer^+^ CD8^+^ T cell responses as well as IFNγ and TNFα responses compared to control mice that received isotype antibody (**Fig 6D-H**). These results suggest that helminth induced IL-10 likely suppresses T cell response to mRNA S6P vaccine.

## DISCUSSION

In this study, we evaluated the impact of intestinal helminths on SARS-CoV-2 spike vaccine responses using a mouse model of hookworm infection. Although intestinal helminth infection did not substantively affect vaccine-induced antibody responses, T cell responses were impacted. The T cell defect was evident regardless of whether Hpb was given prior to the prime or booster vaccine dose, suggesting helminth infection likely interferes at multiple stages of T cell maturation. Hpb infection impaired vaccine-induced T cell responses induced by two distinct vaccine platforms: lipid-encapsulated mRNA (mRNA S6P) and adenoviral-vectored (Ad26.COV2.S) vaccines. Helminth infection did not appear to skew the cytokine profile of antigen-specific T cells towards IL-4 or IL-10 but rather suppressed priming and polyfunctionality of the responses. Challenge experiments showed that Hpb did not substantively affect vaccine-mediated protection against ancestral SARS-CoV-2 strain WA1/2020 D614G but compromised protection in the lungs against the heterologous Omicron variant BA.5.5. This result might be explained by lineage-dependent differences of the immune correlates of Wu-1 spike vaccine protection^43^; whereas WA1/2020 D614G is more readily restricted by serum neutralizing antibodies, which were unaffected by Hpb infection, protection against the antigenically-shifted Omicron variant BA.5.5 likely requires T cells because of evasion of serum neutralizing antibody responses.

Meta-analyses of human data suggest that helminth infection has a greater adverse impact on the immunogenicity live-attenuated vaccines such as measles and Bacille Calmette-Guérin (BCG)^19,20^. In our experiments, T cell responses after immunization with mRNA or adenoviral-vectored vaccines also were negatively impacted by Hpb infection. Hpb had a greater negative effect on T cell responses to mRNA S6P than Ad26COV2.S, which could be due to differences in timing, dosing, or distinct mechanisms of T cell priming. In comparison, we observed smaller changes in B cell or antibody responses, possibly because the type 2 cytokines evoked by Hpb (*e.g*., IL-4) do not hinder B cell responses^44^. It is possible that adenovirus vector-based vaccines may have advantages in helminth-endemic areas, especially in instances where T cell responses are required for protection.

Numerous studies have implicated the STAT6 signaling pathway and type 2 immunity as a dominant mediator by which intestinal helminths suppress immune response^12,38,39^. However, we report that intestinal helminth mediated suppression of T cell responses to mRNA vaccines occurs independently of STAT6 and likely, the upstream cytokines such as IL-4 and IL-3. Instead, based on statistical analysis, IL-10, IL-17, IL-1α, and G-CSF were predictive factors for poor spike-specific CD8^+^ T cell responses. We speculated that IL-10 might be a key regulator because of its immunosuppressive functions in infection and inflammation^45^. Indeed, blocking of IL-10 in helminth infected mice restored vaccine induced CD8^+^ T cell responses. Although IFNγ^+^TNFα^−^ CD8^+^ T cell responses were fully restored, there was only partial restoration of IFNγ^+^TNFα^+^ CD8^+^ T cells, suggesting that additional helminth generated signals might dampen the induction of TNFα expression. Whether these additional signals include other cytokines such as IL-17A, G-CSF or IL-1α remains to be determined. Our findings may be relevant to human responses since children infected with hookworms have elevated levels of IL-10 circulating in their blood^46,47^, and diminished cellular responses to vaccines in helminth-infected individuals has been linked to IL-10^48,49^.

One mechanism by which intestinal helminth infection and IL-10 could affect CD8^+^ T cell responses is by inhibiting maturation of the APCs involved in priming and CD4^+^ T cell help. Indeed, helminths and their products can inhibit expression of MHC class II or co-stimulatory molecules and render DCs less responsive to TLR ligands^50–52^. In some cases, this was associated with increased production and circulation of IL-10^53^. Alternatively, elevated levels of IL-10 could have direct effects on T cells. Consistent this idea, antigen-independent stimulation of T cells with PMA/ionomycin was dampened in helminth infected vaccinated mice. Exposure to IL-10 could raise threshold of signals necessary for production of cytokines by effector T cells^54^. The precise mechanisms by which Hpb induced IL-10 impairs vaccine-induced T cell responses warrants further examination.

The rationale for failure of vaccines to induce optimal immune response in certain populations has largely remain enigmatic. Our data in mice suggests that the elevated levels of IL-10 seen in helminth infected individuals might blunt vaccine-induced T cell responses^48,49^, especially in those given mRNA vaccines. This finding might be particularly relevant as this platform likely will be a future mainstay against infectious and non-infectious diseases including in populations with endemic helminth infection^55,56^.

Vaccine efficacy study results often are not generalizable to all populations. Vaccines that confer immunity to infants or children in developed countries fail to achieve similar results in low income countries^13,14,57^, which have a high prevalence of helminth infection due to limited access to clean water^7^. Similarly, lower than expected efficacy of vaccines against malaria, *Mycobacterium tuberculosis*, rotavirus, and hepatitis B virus in helminth endemic regions suggests that helminth infections may be an important co-factor that negatively affects vaccine immunity^11,19,57^. Our results support the idea that immunization campaigns should consider evaluating the effect of chronic helminth infections on vaccine immunogenicity and outcome.

### Limitations of the study

We acknowledge several limitations to our study. (a) Our experiments showing an adverse effect of enteric helminth infection on SARS-CoV-2 vaccine-induced T cell responses were performed in mice. Studies in helminth-infected human cohorts are required to corroborate these findings. (b) Although we defined an important role of IL-10, whether IL-10 acts directly on T cells or indirectly through APCs was not determined. (c) Although our IL10-reporter mouse studies narrowed down the cellular sources of IL-10 in lymphoid tissues after Hpb infection, the cell type that dominantly contributes to the CD8^+^ T cell phenotype remains unclear. (d) We tested the impact of Hpb infection, which persists principally in the proximal small intestine of mice. Other helminths that occupy distinct intestinal niches or invade tissues could differentially affect vaccine-induced immunity. (e) Our immunization studies did not consider the effects of pre-existing virus infection-induced immunity or hybrid immunity (vaccination and natural infection), which are now common scenarios for SARS-CoV-2.

In summary, our study in mice shows that enteric helminth infection has detrimental effects on T cell responses induced by two SARS-CoV-2 vaccine platforms. Impairment in antigen-specific CD8^+^ T cell responses through an IL-10-dependent signaling axis likely contributed to compromised protection in the lung against a SARS-CoV-2 Omicron variant that escapes neutralization by serum antibodies. Since, one-third of the global population lives in endemic areas, helminths should be considered as key cofactors that can modulate vaccine immunogenicity and efficacy.

## SUPPLEMENTAL FIGURES

**Figure S1. Effect of Hpb infection on antibody and B cell responses in mice immunized with mRNA S6P vaccine, Related to** Figure 1 **and 2**. (**A**) Scheme of experimental design and groups of mice. Cohorts of 6-week-old male C57BL/6J mice were immunized intramuscularly with 0.5 μg of mRNA S6P and boosted with the same dose 21 days later. One cohort of mice received infective Hpb L3 12 days prior to both prime and boost (P/B) doses, whereas the other two groups received Hpb either 12 days prior to prime (P) or 12 days prior to boosting (B). Some mice were kept unvaccinated (no Hpb infection or vaccination). (**B-C**) Fecal samples were collected from the mice at (**B**) day -1 of prime dose or (**C**) day -1 of booster dose, and egg burden was assessed. Two experiments, n = 8 mice per group. (**D-E**) Serum samples were collected at (**D**) day 15 post-prime and (**E**) day 15 post-boost, and IgG binding to Wu-1 spike or RBD was measured by ELISA. Bars indicate geometric mean values. (**F-I**) Splenocytes were isolated at day 15 post-boost and stained for Wu-1 spike-specific germinal center B cells (GC; CD45^+^CD19^+^CD4^-^IgD^low^GL7^+^Fas^+^) and memory B cells (MBC; CD45^+^CD19^+^CD4^-^ IgD^low^GL7^-^Fas^-^TACI^-^CD138^-^) by flow cytometry. (**F**) Gating strategy for spike-specific germinal center (GL7^+^CD38^+^) and memory (GL7^+^CD38^+^TACI^-^CD138^-^) B cells. CD19^+^IgD^low^ B cells negative for germinal center and plasma cell markers were considered memory B cells. Representative flow plots (**G**), absolute numbers of spike-specific GC B cells (**H**) and MBCs (**I**). Bars indicate mean values. (**J-L**) Serum samples were collected at 30 days post-boost, and neutralizing antibody levels were measured with authentic WA1/2020 D614G (**J**), BA.1 (**K**), and BA.5 (**L**) viruses. Bars indicate geometric mean values. (**D-I**) Two experiments, *n* = 8 mice per group, and dotted lines show limit of detection [LOD]. Statistical analysis: (**B, C**) one-way ANOVA with Dunn’s post-test; all groups are compared to naïve. (**D-L**) one-way Kruskal–Wallis ANOVA with Dunn’s post-test. Comparisons are between uninfected mRNA S6P vaccinated groups and Hpb-infected mRNA S6P vaccinated groups (ns, not significant; \**P* < 0.05, \*\**P* < 0.01).

**Figure S2. CD8^+^ T cell response in mice immunized with mRNA S6P and infected with Hpb, Related to** Figure 1. (**A**) Gating strategy for CD8^+^ T cell responses. Spleens were harvested at 15 dpi, and splenocytes were incubated *ex vivo* with VL8 peptide, and stained intracellularly for IFNγ, TNFα, IL-2, IL-4, and IL-10 expression. (**A**) Gating strategy for measuring total percentages and numbers of IFNγ^+^TNFα^-^, IFNγ^+^TNFα^+^ and IL-2^+^IFNγ^+^ cells among CD44^+^ activated CD8^+^ T cells. (**B**) CD44^+^ CD8^+^ T cells were gated on IFNγ^+^ cells and thereafter for TNFα and IL-2 respectively for quantifying proportionality of polyfunctional response. (**C-F**) Representative flow plots, percentages, and cell numbers of IL-4^+^IFNγ^-^ and IL-10^+^IFNγ^-^ cells among CD44^+^ activated CD8^+^ T cells. Two experiments, n = 10 mice per group. Statistical analysis: one-way ANOVA with Holm-Sidak’s post-test; comparisons are between uninfected mRNA S6P vaccinated groups and Hpb-infected mRNA S6P vaccinated and unvaccinated groups (ns, not significant, \**P* < 0.05).

**Figure S3. CD4^+^ T cell response in mice immunized with mRNA S6P and infected with Hpb and T_H_2 immune response induced by Hpb, Related to** Figure 2. (**A-B**) Spleens were harvested at day 15 post-boost, and splenocytes were incubated *ex v*ivo with spike peptide and stained intracellularly for IFNγ, TNFα, and IL-2 (**A**) or IL-4 and IL-10 (**B**). Two experiments, n = 8 to 10 mice per group, (**C-D**) Spleens were harvested at day 15 post-boost, and splenocytes were stained with Live/dead marker, and antibodies to CD45, CD3, CD4, CD44, Foxp3 and GATA-3. Percentage (**C**) and absolute numbers (**D**) of T_H_2 cells (CD4^+^Foxp3^-^ CD44^+^GATA-3^+^) were calculated. Two experiments, n = 8 to 10 mice per group. Statistical analysis: (**A, B**) one-way ANOVA with Holm-Sidak’s post-test; comparisons are between uninfected mRNA S6P vaccinated groups and Hpb-infected mRNA S6P vaccinated and unvaccinated groups; (**D**) Kruskal-Wallis test with Dunn’s post-test; comparisons are between Hpb-infected and other groups (ns, not significant, \**P* < 0.05, \*\**P* < 0.01, \*\*\**P* < 0.001, \*\*\*\**P* < 0.0001).

**Figure S4. CD8^+^ and CD4^+^ T cell response to Hpb-infected mice immunized with Ad26.COV2.S vaccine, Related to** Figures 1 **and 2**. (**A**) Schematic representation of experimental design. Cohorts of C57BL/6J mice were immunized with the Ad26.COV2.S vaccine and boosted 30 days later. Infective Hpb L3 was given 12 days prior to first (prime) dose, and antigen-specific T cell responses were assessed 10 days after boosting. (**B-C**) Splenocytes were isolated at day 10 post-boost, stained with antibodies and the VL8 class I MHC tetramer, and (**C**) the percentages and numbers of tetramer-positive cells were quantified. (**D-E**) Splenocytes were isolated at day 10 post-boost, and tetramer-positive CD8^+^ T cells were analyzed for expression of CX3CR1 and KLRG1. (**E**) The numbers of three subsets (CX3CR1^-^ KLRG1^-^, CX3CR1^+^KLRG1^-^, and CX3CR1^+^KLRG1^+^ cells) were quantified. For naïve and Hpb-only infected mice, cells were pre-gated on total CD8^+^ T cells. n = 5 mice per group. (**F-G**) At day 10 post-boost, splenocytes were stimulated with the VL8 spike peptide, stained intracellularly for IFNγ, TNFα, and IL-2 and analyzed by flow cytometry. (**G**) Numbers of IFNγ^+^TNFα^-^, IFNγ^+^TNFα^+^ and IL-2^+^IFNγ^+^ among CD44^hi^ (activated) CD8^+^ T cells. (**H-I**) At day 10 post-boost, splenocytes were stimulated with VL8 peptide, stained intracellularly for IFNγ, IL-4 and IL-10 and analyzed. Representative flow cytometry plots of IFNγ and IL-4 (**H**) and IFNγ and IL-10 (**I**) are shown. (**J-L**) Splenocytes were isolated at day 10 post-boost and stimulated *ex vivo* with immunodominant CD4 spike peptide, stained intracellularly for IFNγ and TNFα, and CD4^+^ T cells, and analyzed by flow cytometry. Representative flow cytometry plots (**J**), percentages and numbers (**K**) of IFNγ^+^TNFα^+^ double-positive cells. (**L-M**) Splenocytes were stained for expression of IFNγ, IL-4, and IL-10, and analyzed by flow cytometry and representative flow plots are shown. Two experiments, n = 8 to 10 mice per group. Statistical analysis: one-way ANOVA with Holm-Sidak’s post-test; comparisons are between uninfected and Hpb-infected Ad26.COV2.S immunized groups or between unvaccinated and Ad26.COV2.S immunized groups (ns, not significant, \**P* < 0.05, \*\**P* < 0.01, \*\*\**P* < 0.001, \*\*\*\**P* < 0.0001).

**Figure S5. Egg burden and antigen-specific CD8^+^ T cell response in K18-hACE2 mice immunized with mRNA S6P and infected with Hpb. Related to Figure 3**. (**A**) Cohorts of 6-week-old K18-hACE2 transgenic mice were orally gavaged with Hpb and 12 days later vaccinated with two dose regimen of mRNA S6P (0.5 μg). Four to five weeks post boost, mice were challenged intranasally (IN) with 10^4^ FFU of WA1/2020 D614G or BA.5.5 SARS-CoV-2 variants. (**B**) Fecal samples were collected 11 days after Hpb inoculation of K18-hACE2 and wild-type C57BL/6J mice, and egg burden was measured. Two experiments, n = 8 mice per group, (**C**) Spleens were harvested at day 15 post-boost in K18-hACE2 transgenic mice, and splenocytes were stained with live/dead marker, CD45, CD3, CD8, CD44 and VL8 tetramer. Representative flow cytometry plots of tetramer positive CD8^+^ T cells with numbers of cells measured. n = 6 mice per group. Mann-Whitney test (ns = not significant, \*\**P* < 0.01).

**Figure S6. Hpb infection makes vaccine induced CD8^+^ T cell hyporesponsive to antigen-dependent and independent stimulation, Related to Figure 4**. (**A-B**) Hpb infected mice were vaccinated with two dose regimen of mRNA S6P and at day 10 post-boost, splenocytes were stimulated *ex vivo* with either VL8 peptide, CD3/CD28 and PMA/ionomycin, and stained intracellularly with IFNγ and TNFα. Representative flow plots shown (**A**) and numbers (**B**) of IFNγ^+^TNFα^-^ single-positive cells and IFNγ^+^TNFα^+^ double-positive cells were quantified. (**A-B**) Two experiments, n = 8 mice per group. Statistical analysis: one-way ANOVA with Holm-Sidak’s post-test; comparisons are between mRNA S6P immunized group (grey) with all other groups (ns = not significant, \**P* < 0.05, \*\**P* < 0.01, \*\*\**P* < 0.001, \*\*\*\**P* < 0.0001).

**Figure S7. Serum cytokine analysis in Hpb infected and uninfected mRNA S6P immunized mice, Related to** Figure 5. (**A-B**) Spearman correlation analysis of serum cytokine clusters between uninfected mice vaccinated with mRNA S6P (grey dots) and Hpb-infected vaccinated mice (green dots) at day 1 (**A**) and 3 post-prime (**B**) compared to total numbers of splenic VL8 tetramer positive CD8^+^ T cells at day 15 post-boost. Statistical analysis: (**A, B**) Mann-Whitney test; Clustering analysis of cytokines levels was performed with K-means clustering method (see Methods; n = 6 per group). Non-parametric Spearman correlation analysis was performed using a 95% confidence interval (ns, not significant, ** *P* < 0.01, *** *P* < 0.001). (**C**) Serum samples were collected from Hpb infected WT and Hpb infected *Stat6*^-/-^ mice at day 3 post-prime and cytokines IL-4, IL-5, IL-13, ST2, G-CSF, IL-17A, IL-10 and IL-1α were measured and represented as Log_2_ fold change *Stat6*^-/-^ mice compared to WT mice. Two experiments, n = 6 mice per group.

**Figure S8. Gating strategy of various immune cell populations in the spleen, Related to** Figure 6. 10BiT (IL-10 reporter) mice were infected with Hpb and 12 days later immunized with mRNA S6P. Splenocytes were harvested at day 3 post immunization and stained with surface markers and analyzed by flow cytometry.

## Acknowledgements.

This study was supported by the NIH (R01 AI157155, NIAID Centers of Excellence for Influenza Research and Response (CEIRR) contracts 75N93021C00014 and 75N93019C00051, to M.S.D.). We thank Adrianus Boon for providing Ad26.COV2.S vaccine and Mehul Suthar for the BA.5 isolates used in this study. We thank Larry Yang and Diane Bender from Bursky Center for Human Immunology and Immunotherapy Programs (CHIIPS) at Washington University for help with tetramers and cytokine assays.

## Author contributions

P.D. and C.E.K. performed helminth infections, immunizations, binding assays, challenge experiments, viral burden assays, and flow cytometry experiments. P.D. performed and analyzed most of the T cell experiments. B.Y. performed neutralization assays. C.Y.L. performed B cell staining. T.G.S., A.C.S., F.T.C., S.P.R., and R.P.S., performed serum cytokine analysis and correlation analysis. J.F.U provided initial batch of helminth parasites. D.K.E. and S.M.E. provided mRNA vaccines and helped in the design of vaccination experiments. P.D., L.B.T., R.P.S., and M.S.D. designed studies. P.D. and M.S.D wrote the initial draft, with the other authors providing editorial comments.

## STAR METHODS

### RESOURCE AVAILABILITY

#### Lead Contact

Further information and requests for resources and reagents should be directed to the Lead Contact, Michael S. Diamond.

#### Materials Availability

All requests for resources and reagents should be directed to and will be fulfilled by the Lead Contact author. This includes mice, antibodies, viruses, and helminths. All reagents will be made available on request after completion of a Materials Transfer Agreement. Preclinical mRNA vaccines can be obtained under an MTA with Moderna (contact: Darin Edwards).

#### Data and code availability

All data supporting the findings of this study are available within the paper and are available from the corresponding author upon request.

### EXPERIMENTAL MODEL AND SUBJECT DETAILS

#### Mouse experiments

All mice studies were conducted in strict accordance with the recommendations of the Guide for the Care and Use of Laboratory Animals of the National Institutes of Health. Animal experiments were performed as specified in protocols approved by the Institutional Animal Care and Use Committee (IACUC) at the Washington University School of Medicine (Assurance Number: A3381-01). All dissections and virus inoculations were performed under anesthesia, induced and maintained by using ketamine hydrochloride and xylazine, and every effort was made to minimize suffering. Wild-type male C57BL/6J mice (cat no. 000664) were obtained from The Jackson Laboratory and used at 6 to 8 weeks of age for all experiments. Heterozygous K18-hACE2 C57BL/6J mice (strain: 2B6.Cg-Tg(K18-ACE2)2Prlmn/J, cat no. 34860) were obtained from The Jackson Laboratory. Stat6^tm1Gru^ (*Stat6*^-/-^) (005977) were purchased from Jackson Laboratories and bred at Washington University under specific pathogen-free conditions. 10BiT reporter mice crossed to Foxp3^gfp^ reporter mice were kindly provided by Chyi Song Hsieh (Washington University in St. Louis)^42^. All mice were housed in the animal facility, fed standard chow diets, and kept at ∼70°F with humidity at ∼50%, and the dark/light cycle was 12 h/12 h.

#### Cells

African green monkey Vero-TMPRSS2 ^58^ and Vero-hACE2-TMPRRS2 cells ^59^ were cultured at 37°C in DMEM supplemented with 10% FBS, 10 mM HEPES pH 7.3, and 100LULper mL of penicillin–streptomycin. Vero-TMPRSS2 cells were supplemented with 5 µg perLmL of blasticidin. Vero-hACE2-TMPRSS2 cells were supplemented with 10LµgLperLmL of puromycin. All cells routinely tested negative for mycoplasma using a polymerase chain reaction (PCR)-based assay.

#### Viruses

WA1/2020 D614G virus was described previously^60,61^. BA.5 (hCoV-19/USA/CA-Stanford-79_S31/2022) and BA5.5 (hCoV-19/USA/COR-22-063113/2022) isolates were generously provided as a gift by M. Suthar (Emory University) and A. Pekosz (Johns Hopkins School of Medicine), respectively. All viruses were passaged once on Vero-TMPRSS2 cells and subjected to next-generation sequencing to confirm the introduction and stability of substitutions^59^. All virus experiments were performed in an approved biosafety level 3 (BSL-3) facility at Washington University School of Medicine.

#### Viral antigens

Recombinant soluble S and RBD proteins from Wuhan-1 SARS-CoV-2 strains were expressed as described^32^. Recombinant proteins were produced in Expi293F cells (ThermoFisher) by transfection of DNA using the ExpiFectamine 293 Transfection Kit (ThermoFisher). Supernatants were harvested 3 days post-transfection, and recombinant proteins were purified using Ni-NTA agarose (ThermoFisher), then buffer-exchanged into PBS and concentrated using Amicon Ultracel centrifugal filters (EMD Millipore).

### METHOD DETAILS

#### Preclinical mRNA vaccine mRNA S6P and lipid nanoparticle production process

A sequence-optimized mRNA encoding prefusion-stabilized Wuhan-Hu-1 (mRNA S6P) SARS-CoV-2 S-6P (hexaproline-stabilized) protein was synthesized *in vitro* using an optimized T7 RNA polymerase-mediated transcription reaction with complete replacement of uridine by N1m-pseudouridine ^62^. A non-translating control mRNA was synthesized and formulated into lipid nanoparticles as previously described ^63^. The reaction included a DNA template containing the immunogen open-reading frame flanked by 5’ untranslated region (UTR) and 3’ UTR sequences, and was terminated by an encoded polyA tail. After RNA transcription, the cap-1 structure was added using the vaccinia virus capping enzyme and 2L-*O*-methyltransferase (New England Biolabs). The mRNA was purified and kept frozen at –80°C until further use.

The mRNA was encapsulated in a lipid nanoparticle through a modified ethanol-drop nanoprecipitation process described previously^64^. Ionizable, structural, helper, and polyethylene glycol lipids were briefly mixed with mRNA in an acetate buffer, pH 5.0, at a ratio of 2.5:1 (lipid:mRNA). The mixture was neutralized with Tris-HCl, pH 7.5, sucrose was added as a cryoprotectant, and the final solution was sterile-filtered. Vials were filled with formulated lipid nanoparticle and stored frozen at –80°C until further use.

#### ELISA

Purified recombinant Wuhan-1 spike or RBD proteins were coated onto 96-well Maxisorp clear plates at 2 μg/mL (spike) or 2 μg/mL (RBD) in 50 mM Na_2_CO_3_ pH 9.6 (50 μL) overnight at 4°C. Coating buffers were aspirated, and wells were blocked with 200 μL of 1X PBS + 0.05% Tween-20 + 2% BSA + 0.02% NaN_3_ (Blocking buffer, PBSTBA) overnight at 4°C. Sera were serially diluted in blocking buffer and added to the plates. Plates were incubated for 1 h at room temperature and then washed 3 times with PBST, followed by addition of 50 μL of 1:2000 anti-mouse IgG-HRP (Southern Biotech Cat. #1030-05) in PBST. Following a 1 h incubation at room temperature, plates were washed 3 times with PBST and 50 μL of 1-Step Ultra TMB-ELISA was added (ThermoFisher Cat. #34028). Following a 5 to 10-min incubation, reactions were stopped with 50 μL of 2 M H_2_SO_4_. The absorbance of each well at 450 nm was determined using a microplate reader (BioTek) within 5 min of addition of sulfuric acid. The endpoint serum dilution was calculated with curve fit analysis of optical density (OD) values for serially diluted sera with a cut-off value set to mean plus six times the standard deviation of the background signal. Optical density (OD) measurements were taken at 450 nm, and titers were determined using a 4-parameter logistic curve fit in Prism v.8 (GraphPad 112 Software, Inc.) and defined as the reciprocal dilution at approximately optical density 450 of 1 (normalized to a mouse standard on each plate).

#### Focus reduction neutralization test

Serial dilutions of serum samples were incubated with 10^2^ focus-forming units (FFU) of SARS-CoV-2 strains WA1/2020 D614G, BA.1, BA.5 for 1 hour at 37°C. Antibody-virus complexes were added to Vero-TMPRSS2 cell monolayers in 96-well plates and incubated at 37°C for 1 hour. Subsequently, cells were overlaid with 1% (w/v) methylcellulose in Eagle’s minimal essential medium (Thermo Fisher Scientific). Plates were harvested 30 hours later by removing overlays and fixed with 4% paraformaldehyde in PBS for 20 min at room temperature. Plates were washed and sequentially incubated with an oligoclonal pool of SARS2-2, SARS2-11, SARS2-16, SARS2-31, SARS2-38, SARS2-57, and SARS2-71 ^65^ anti-spike protein antibodies and HRP-conjugated goat anti-mouse IgG (Sigma-Aldrich, catalog no. A8924, RRID: AB_258426) in PBS supplemented with 0.1% saponin and 0.1% bovine serum albumin. SARS-CoV-2–infected cell foci were visualized using TrueBlue peroxidase substrate (KPL) and quantitated on an ImmunoSpot microanalyzer (Cellular Technologies).

#### Helminth inoculations and egg burden quantification

Infective Hpb L3 were generated as described^66^. Hpb viability was checked by microscope, and their numbers were quantified before use. Mice were gavaged with 300 L3 Hpb before prime and/or 150 L3 Hpb before boost or PBS (control) using 20-gauge x 38 mm plastic feeding tubes (Instech; FTP-20-38). Twelve days after Hpb infection, mice were immunized and boosted intramuscularly with 0.5 μg of mRNA S6P in 50 μL volume. Some mice received Hpb before prime and boost whereas other groups received either before prime or before boost. Fresh fecal pellets were collected at indicated time points and eggs per gram of feces were enumerated as a readout of worm fitness. Briefly, two to three fecal pellets were collected from each mouse, weighed, and homogenized in 1 mL of PBS. Homogenized stool was added to tubes containing 1 mL of PBS and 2 mL of saturated NaCl solution. Tubes were mixed by shaking and allowed to settle for 5 to 10 min for the eggs to float on the top. Using a sterile pipette, 540 μL of the mixed solution was loaded onto McMaster 2-chamber egg counter. The worm eggs were counted within the grid and the eggs per gram feces were determined by the following formula: eggs/g = egg counts x 26.67 weight of feces (g). The 26.67 in the formula is obtained from the total volume in which the fecal pellets are suspended (4 mL) divided by the volume within the grid (0.15 mL).

#### Immunization and viral challenge

Helminth inoculated or naïve mice were immunized and boosted with mRNA S6P intramuscularly at 0.5 μg dose in a 50 μl volume. Animals were bled 15 days after prime or boosting for immunogenicity analysis of sera. Spleens were harvested at day 15 post-boost for flow cytometry analysis. At day 30 post-boost, mice were bled and then challenged one week later with 50 μL of 10^4^ FFU of WA1/202 DG14G or BA.5.5 by intranasal administration. Lungs, nasal turbinates, and nasal washes (collected in 500 μL of 0.5% bovine serum albumin in phosphate buffered saline) were harvested five days after inoculation for virological analysis. For adenoviral vaccine experiments, mice were immunized intramuscularly in both legs with thawed Ad26COV2.S (total of 5 x 10^9^ viral particles) and boosted with the same dose 30 days later.

#### Measurement of viral burden

Tissues were weighed and homogenized with zirconia beads in a MagNA Lyser instrument (Roche Life Science) in 1 ml of DMEM medium supplemented with 2% heat-inactivated FBS. Tissue homogenates were clarified by centrifugation at 9,600 x g for 5 min and extracted using the MagMax mirVana Total RNA isolation kit (Thermo Fisher Scientific) on the Kingfisher Flex extraction robot (Thermo Fisher Scientific). RNA was reverse transcribed and amplified using the TaqMan RNA-to-CT 1-Step Kit (Thermo Fisher Scientific). Reverse transcription was carried out at 48°C for 15 min followed by 2 min at 95°C. Amplification was accomplished over 50 cycles as follows: 95°C for 15 s and 60°C for 1 min. Copies of SARS-CoV-2 *N* gene RNA in samples were determined using a published assay^67^.

#### Viral plaque assay

Titration of infectious SARS-CoV-2 was performed as previously described. Briefly, lung homogenates were serially diluted and added to Vero-TMPRSS2-hACE2 cell monolayers in 24-well tissue culture plates for 1 h at 37°C. Cells were then overlaid with 1% (w/v) methylcellulose in MEM with 2% FBS and incubated for 24 h (WA1/2020 N501Y/D614G) or 72 h (BA.5.5). Subsequently, cells were fixed with 4% paraformaldehyde in PBS for 20 min at room temperature before staining with 0.05% (w/v) crystal violet in 20% methanol. Viral plaques were counted manually.

#### T cell analysis

Spleens were harvested at day 15 after boosting for mRNA S6P studies and day 10 after boosting for Ad26COV2.S experiments, and single cell suspensions were generated after tissue disruption and passage through a 70-μm cell strainer. Splenocytes were pelleted by centrifugation, and erythrocytes were lysed using ACK lysis buffer (Thermo Fisher). Cells then were re-suspended in RPMI 1640 media supplemented with 10% FBS, 1% HEPES, 1% L-glutamine, and 0.1% β-mercaptoethanol. For peptide stimulation, splenocytes were incubated separately with class I MHC (peptide sequence S_262-270_: AAAYYVGYL), (VL8, peptide sequence S_539-546_: VNFNFNGL) (1 μg/mL), or class II MHC (peptide sequence S_62-76_: VTWFHAIHVSGTNGT) spike peptides (4 μg/mL) overnight at 37°C in the presence of Brefeldin A (1:500, Invitrogen). The following day, cells were washed and stained with Fc block (Clone 93; Cat: 101320; BioLegend), CD45 (BUV395; Clone 30-F11; Cat: 564279; BD Biosciences), CD8β (PerCP/Cy5.5; Clone YTS156.7.7; Cat: 126610; BioLegend), CD4 (FITC; Clone GK1.5; Cat: 100406; BioLegend), CD44 (APC/Cy7; Clone IM7; Cat: 103028; BioLegend) for 30 min at 4°C in FACS buffer (1x PBS with 2% FBS and 2 mM EDTA). Dead cells were excluded using Live/Dead (Thermo Fisher) that was added concurrently with staining. Following this, cells were washed, fixed with FoxP3 transcription factor permeabilization kit (eBiosciences; Cat: 00-5523-00), and stained for intracellular IFN-γ APC; Clone XMG1.2; Cat: 505810; BioLegend) and TNF-α (PE/Cy7; Clone MP6-XT22; Cat: 25-7321-82; Invitrogen), IL-2 (PE; Clone JES6-5H4; Cat: 503808; BioLegend), IL-4 (PE-Cy7; Clone 11B11; Cat: 25-7041-82; Invitrogen) and IL-10 (PE; Clone JES-16E3; Cat: 505008; BioLegend) using BD fixation/permeabilization kit (BD Biosciences) according to the manufacturer’s instructions. Cells were processed by flow cytometry on a Cytek Aurora and analyzed using FlowJo software version 10.4.2).

For CD3/CD28 stimulation, 96-well flat-bottom plates were incubated overnight at 4°C with anti-CD3 (Clone 17A2; Cat: 16-0032-82; ThermoFisher) at 2 μg/mL in 1X PBS. Next day, excess anti-CD3 was flicked off from the plates and splenocytes mixed with anti-CD28 (Clone 37.51; Cat: 16-0281-82; ThermoFisher) were added and incubated for 8 hours at 37°C. For PMA/ionomycin stimulation, Cell Stimulation Cocktail (500X; Cat: 00-4970-03; eBiosciences) was mixed with splenocytes at 2 μl/mL final concentration according to the manufacturer’s protocol and incubated for 8 hours at 37°C. Cells were washed and stained with fluorophore conjugated antibodies as described earlier.

For tetramer staining, spleen cells were plated on 96-well round bottom plates, washed, and surface stained with fluorophore-conjugated antibodies in FACS buffer (1x PBS, 2% FCS and 2 mM EDTA). Cells stained with Fc block (Clone 93; Cat: 101320; BioLegend), CD45 (BUV395; Clone 30-F11; Cat: 564279; BD Biosciences), CD3 (BV711; Clone 145-2C11; Cat: 563123; BD Biosciences), CD8α (PerCP/Cy5.5; Clone 53-6.7; Cat: 100734; BioLegend), CD4 (BV785; Clone GK1.5; Cat: 100453; BioLegend), CD44 (APC/Cy7; Clone IM7; Cat: 103028; BioLegend), VL8 tetramer-PE conjugated, CX3CR1 (APC; Clone SA011F11; Cat: 149007; BioLegend), KLRG1 (PE-Cy7; Clone 2F1; Cat: 25-5893-80, eBioscience) for 60 min at room temperature. For immune cell staining in IL-10BiT mice, two staining panels were made a) T and NK cell panel: Fc block (Clone 93; Cat: 101320; BioLegend), CD45 (BUV395; Clone 30-F11; Cat: 564279; BD Biosciences), CD3 (BV711; Clone 145-2C11; Cat: 563123; BD Biosciences), CD8α (PerCP/Cy5.5; Clone 53-6.7; Cat: 100734; BioLegend), CD4 (BV785; Clone GK1.5; Cat: 100453; BioLegend), NK1.1 (PE-Cy7; Clone PK136; Cat: 108714; BioLegend) CD90.1/Thy-1.1 (APC; Clone OX-7; Cat: 202526; BioLegend) and b) B cell and innate cell panel: Fc block (Clone 93; Cat: 101320; BioLegend), CD45 (BUV395; Clone 30-F11; Cat: 564279; BD Biosciences), CD3 (BV711; Clone 145-2C11; Cat: 563123; BD Biosciences), B220 (BV650; Clone RA3-6B2; Cat: 103241; BioLegend), CD11b (PerCP/Cy5.5; Clone M1/70; Cat: 101228; BioLegend), Gr-1 (BV421; Clone RB6-8C5; Cat: 108445; BioLegend), F4/80 (APC/Cy7; Clone BM8; Cat: 123118; BioLegend), CD11c (PE-Cy7; Clone HL3; Cat: 558079; BD Biosciences), MHC-II (AF700; Clone M5/114.15.2; Cat: 107622; BioLegend), XCR1 (PE; Clone ZET; Cat: 148204; BioLegend), CD90.1/Thy-1.1 (APC; Clone OX-7; Cat: 202526; BioLegend). Dead cells were excluded using Live/Dead (Thermo Fisher) staining. Cells were incubated with antibodies for 30 minutes at 4°C. Cells were fixed with 2% paraformaldehyde for 5 min at room temperature, washed with FACS buffer and re-suspended in FACS buffer. Samples were processed by flow cytometry on a Cytek Aurora and analyzed using FlowJo software version 10.4.2).

#### Cytokine Luminex assay

Blood samples were collected in microtainer blood collection tubes (BD 365967), centrifuged and serum was transferred into separate tubes and kept frozen. Serum samples were thawed and used in 48-Plex Mouse ProcartaPlex^TM^ Panel (EPX480-20834-901) and used according to manufacturer’s recommendation.

#### Lung histology

Lungs of euthanized mice were inflated with ∼2 ml of 10% neutral buffered formalin using a 3-ml syringe and catheter inserted into the trachea and kept in fixative for 7 days. Tissues were embedded in paraffin, and sections were stained with hematoxylin and eosin. Images were captured using the NanoZoomer (Hamamatsu) at the Alafi Neuroimaging Core at Washington University.

### QUANTIFICATION AND STATISTICAL ANALYSIS

Statistical significance was assigned when *P* values were < 0.05 using GraphPad Prism version 9.3. Tests, number of animals, median values, and statistical comparison groups are indicated in the Figure legends. Changes in infectious virus titer, viral RNA levels, or serum antibody responses were analyzed by one-way ANOVA with a post-test correction when comparing three or more groups. When comparing two groups, a Mann-Whitney test was performed. Best-fit lines were calculated using non-linear regression analyses. For serum cytokine assay, statistical analyses were conducted by using R (version 4.2.3) and GraphPad Prism software (version 9.3.1 and 9.5.1). Clustering of cytokines levels was performed with K-means clustering method using the optimal number of clusters identified by gap statistics ^68^. Cluster centers were compared using a Mann-Whitney test. For feature selection, scaled values of significantly different cytokines (p-value < 0.05) were used as input for a Lasso regression (glmnet R package -v 4.1-7), and cytokines with non-zero coefficients were selected. Nonparametric Spearman correlation analysis was performed using a 95% confidence interval.

