## Supplementary figures and images for "Intestinal helminth infection impairs vaccine-induced T cell responses and protection against SARS-CoV-2"

### Figure S1

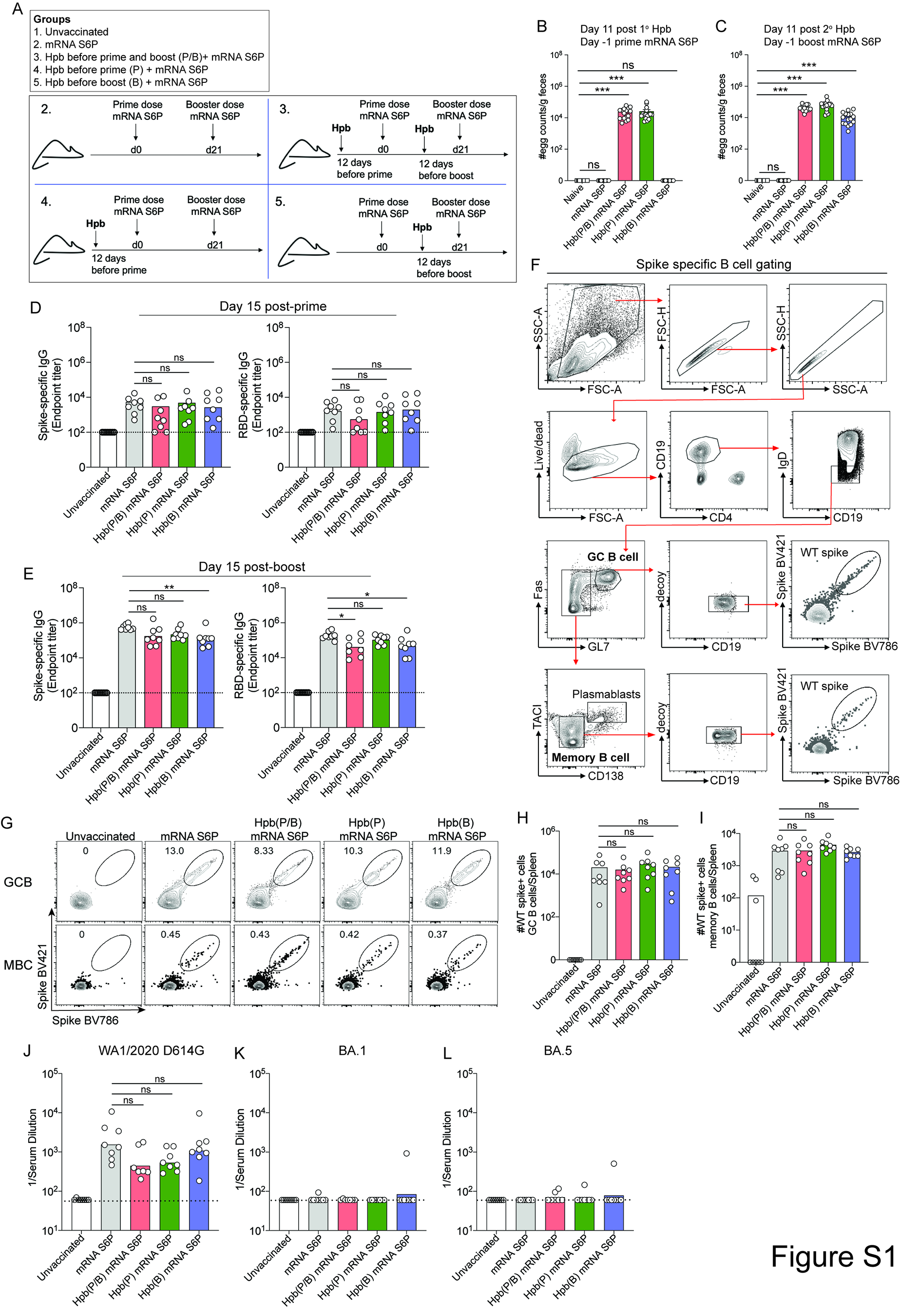

### Figure S2

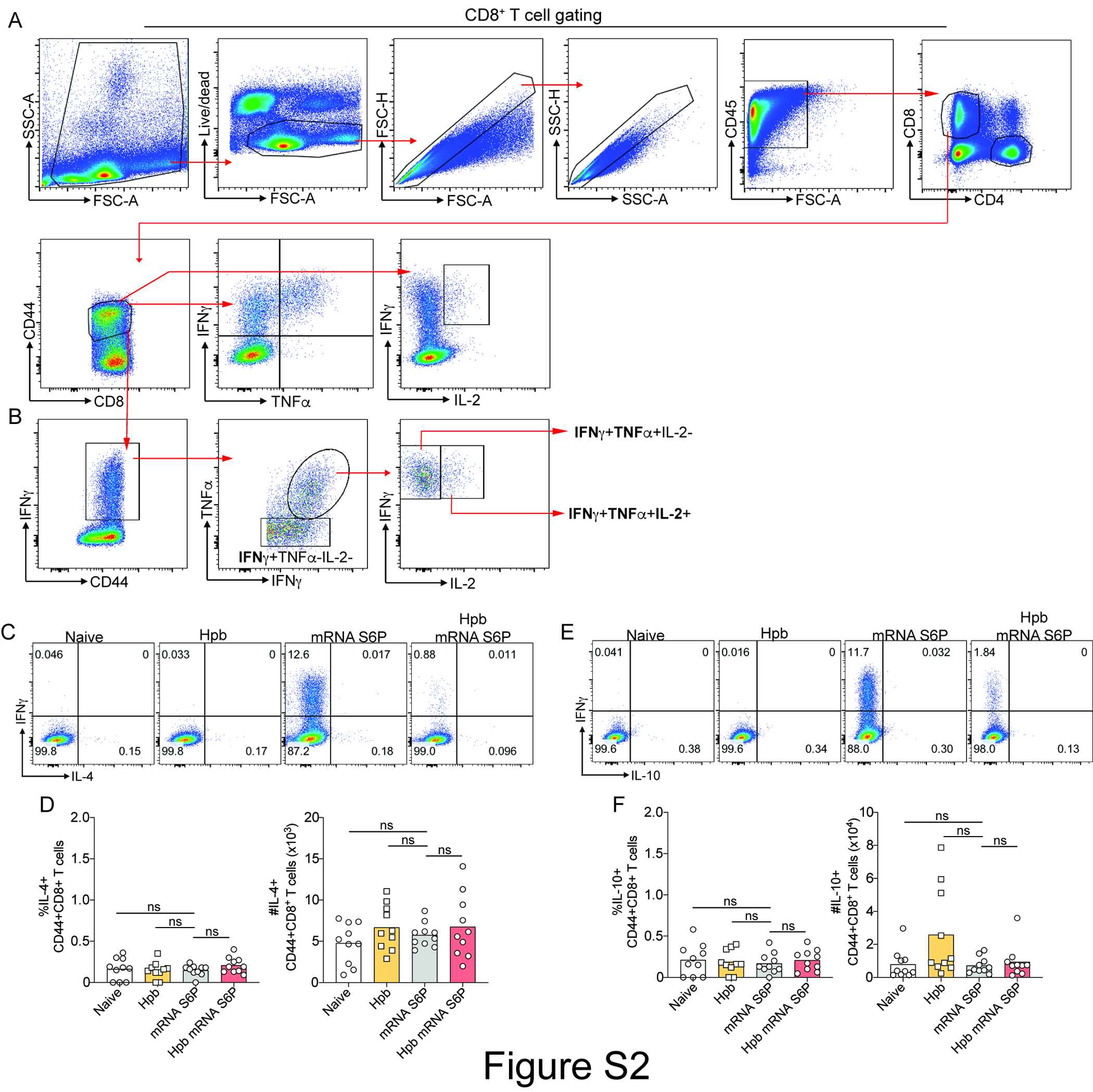

### Figure S3

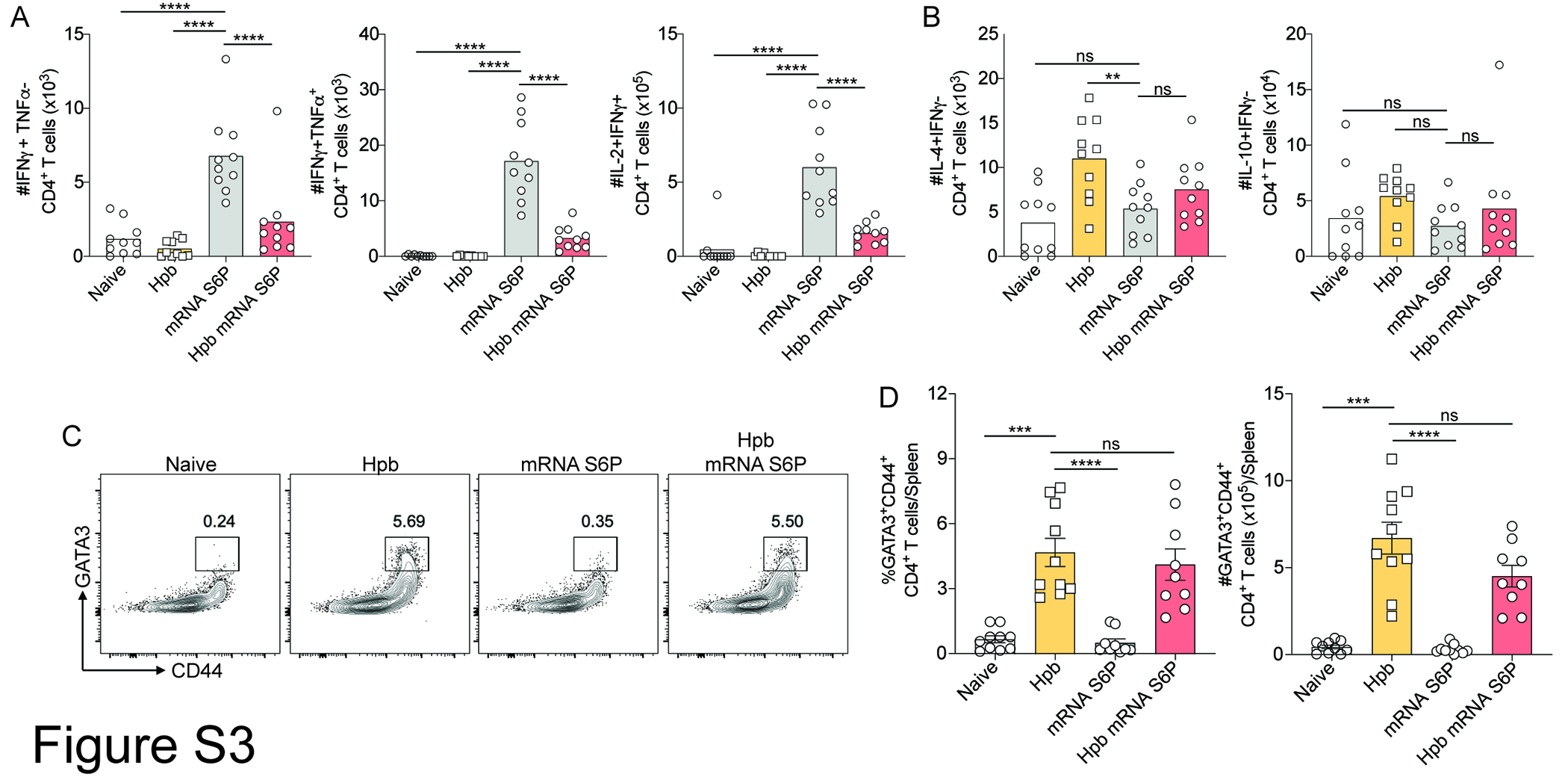

### Figure S4

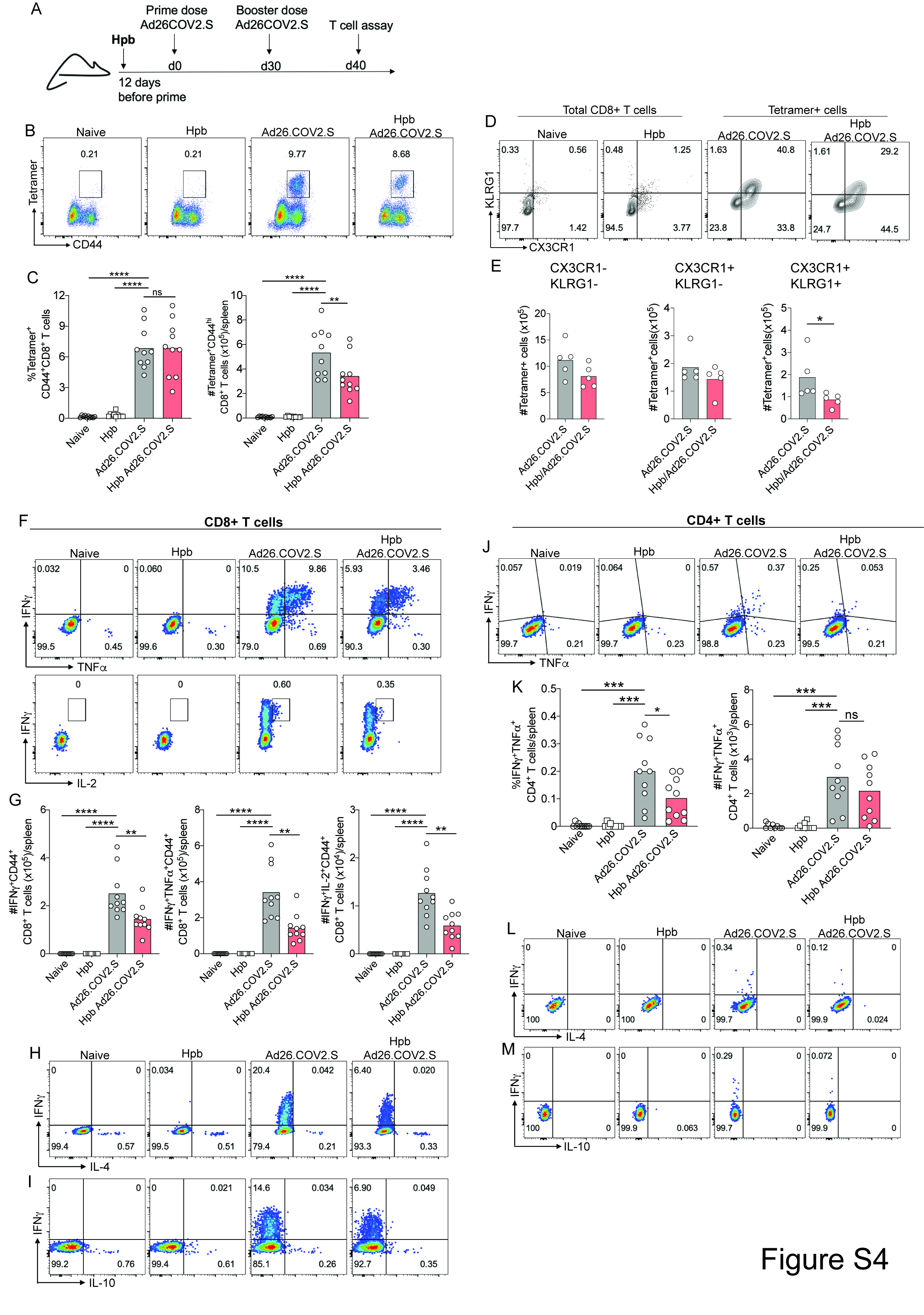

### Figure S5

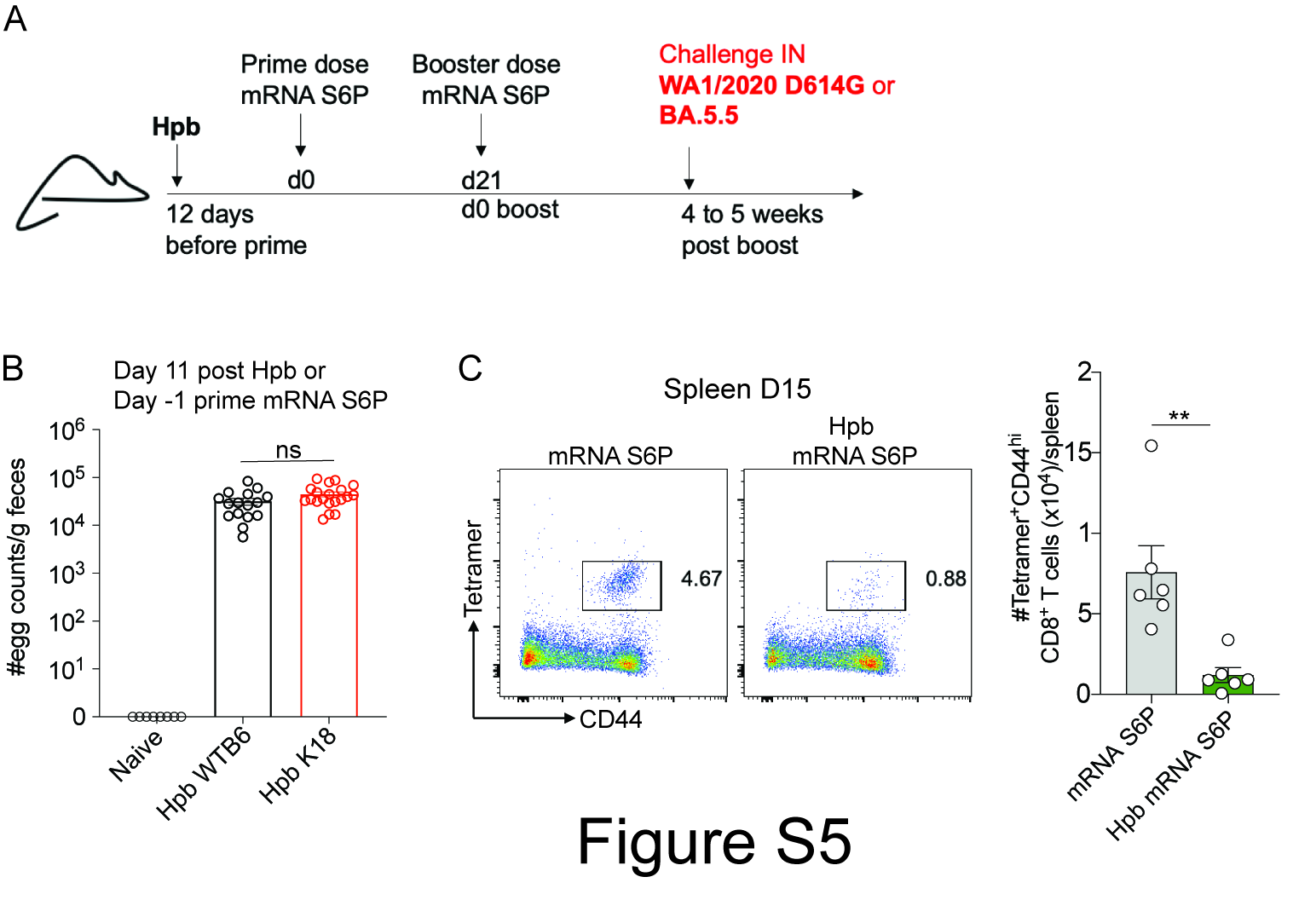

### Figure S6

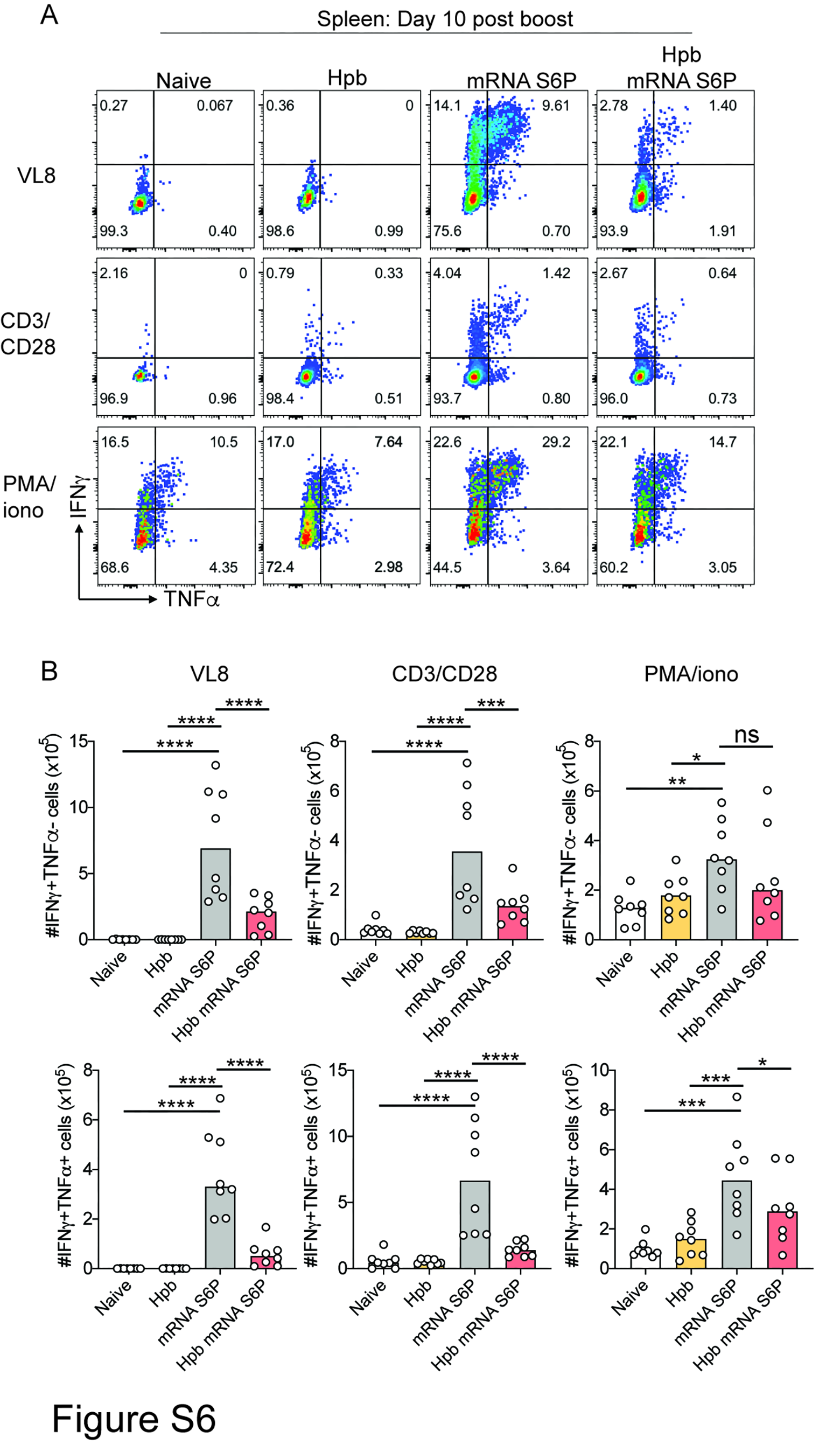

### Figure S7

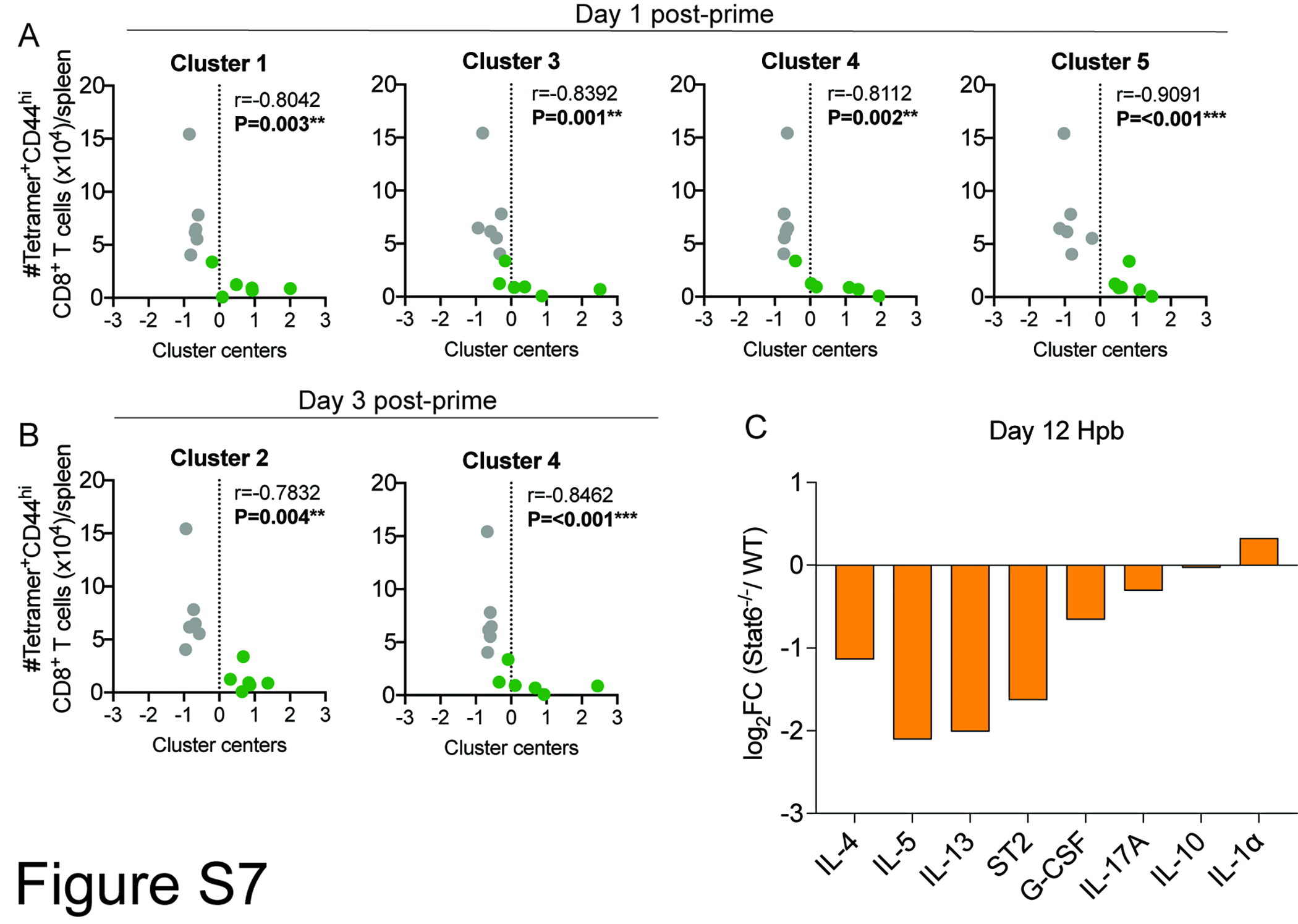

### Figure S8

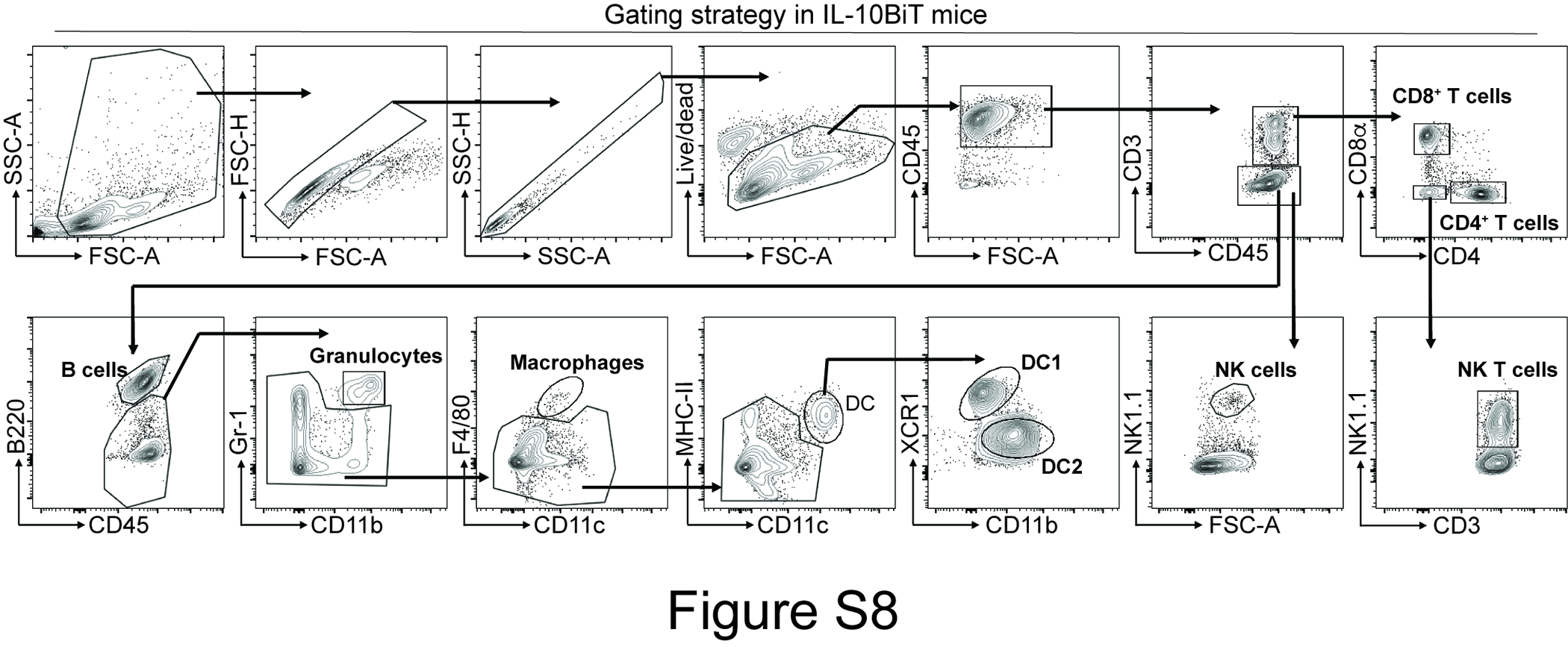
